# Systematically investigating and identifying bacteriocins in the human gut microbiome

**DOI:** 10.1101/2024.07.15.603639

**Authors:** Dengwei Zhang, Yinai Zou, Yuqi Shi, Junliang Zhang, Jing Liu, Gengfan Wu, Jian Zhang, Ying Gao, Muxuan Chen, Yong-Xin Li

**Affiliations:** Department of Chemistry and The Swire Institute of Marine Science, The University of Hong Kong, Pokfulam Road, Hong Kong, China; Microbiome Medicine Center, Department of Laboratory Medicine, ZhuJiang Hospital, Southern Medical University, Guangzhou 510280, China

**Keywords:** Bacteriocin, Antimicrobial peptides, Natural products, Antimicrobial activity, Human microbiome, Genome mining, Metagenomics, Omics analysis

## Abstract

Human gut microbiota produces unmodified bacteriocins, natural antimicrobial peptides that protect against pathogens and regulate host physiology. However, current bioinformatic tools limit the comprehensive investigation of bacteriocins’ biosynthesis, obstructing research into their biological functions. Here, we introduce IIBacFinder, a superior analysis pipeline for identifying unmodified class II bacteriocins. Through large-scale bioinformatic analysis and experimental validation, we demonstrate their widespread distribution across the bacterial kingdom, with being niche-specific and enriched in the human microbiome. Analyzing over 280,000 bacterial genomes, we reveal the diverse potential of human gut bacteria to produce these bacteriocins. Guided by meta-omics analysis, we synthesized 26 hypothetical bacteriocins from gut commensal species, with 16 showing antibacterial activities. Further *ex vivo* tests show minimal impact of narrow-spectrum bacteriocins on human fecal microbiota. Our study highlights the huge biosynthetic potential of unmodified bacteriocins in the human gut, paving the way for understanding their biological functions and health implications.

## Introduction

Bacteria are ubiquitously present on Earth, often found in a communal fashion where each species will compete for limited nutrients and space^1^. These common antagonistic interactions are crucial not only to influence the composition of their members but also to maintain the community stability^2,3^. The interference competition can be governed by various types of molecules, such as polyketides^4^, protein toxins^5^, non-ribosomally synthesized peptides (NRPs)^6^, and ribosomally synthesized antimicrobial peptides that are referred to as bacteriocins^7,8^. Despite exhibiting incredible sequence and structural diversity, bacteriocins are usually categorized into two major classes: those that undergo posttranslational modifications by enzymatic tailoring (class I bacteriocins) and unmodified peptides (class II bacteriocins)^9^. On the basis of posttranslational modifications, class I bacteriocins can be further grouped into more than 40 subgroups such as lanthipeptide and lasso peptide, whereas class II bacteriocins are generally divided into four subgroups (class IIa, YGNG-motif containing bacteriocins; class IIb, two-peptide bacteriocins; class IIc, leaderless bacteriocins; class IId, other linear bacteriocins)^10^.

Due to its antimicrobial nature, bacteriocins have been regarded as promising antibiotic alternatives or adjuncts to contain the antimicrobial resistance threats of current antibiotics^7^. Compared to class I bacteriocins, unmodified class II bacteriocins are simpler for their production in heterologous systems and chemical synthesis, thereby rendering it more practical for antimicrobial discovery at a large scale. Although a few class II bacteriocins have been identified generally via traditional isolation techniques (Table S1), they exhibit a variety of biological activities, such as antifungal and antiviral activities^11,12^. This makes class II bacteriocins promising candidates for the discovery of antimicrobials. Additionally, recent studies, albeit limited, have broadened the functions of bacteriocins beyond their antimicrobial nature, suggesting a role in interacting with the host^13–15^. Our recent study of antimicrobial peptides from human microbiota further suggests the largely unexplored potential of bacteriocins in the gastrointestinal tract (GIT) and their potential physiological roles in human health^16,17^. This further emphasizes the significance of studying host-associated class II bacteriocins, particularly those from the GIT where bacteriocins are crucial to human health^18,19^.

In the GIT, bacteriocins act as essential protectors impacting human health. For example, plantaricin EF, a class IIb bacteriocin, can maintain the integrity of the epithelial barrier, reducing the effects of obesity-inducing diets in mice on a high-fat diet^15^. Another instance demonstrated the immunomodulatory activity of a lactococcin 972-like class IId bacteriocin from *Bifidobacterium longum* subsp. *infantis*^20^, a bacterium found in the infant gut. Given that the human microbiome contains 150 times more genes than the human genome^21^, class II bacteriocins within the human microbiome represent an untapped treasure for antimicrobial discovery and play a crucial role in mediating human health. Although previous investigations on hundreds of human GIT reference genomes or human metagenomic data revealed limited biosynthetic potential of class II bacteriocin in human gut microbiome^22,23^, the overall understanding is significantly incomplete, considering over 280,000 reference genomes reconstructed from the human gut microbiome^24^. It is imperative to conduct a comprehensive investigation into class II bacteriocins in the human gut microbiome to better understand their biosynthetic potential, chemical diversity, distribution, and possible biological functions.

In the meta-omics era, with tremendous genomic data sets available, bacteriocin discovery has evolved from a traditional activity screening to a more prominent emphasis on genomics-guided discovery via predicting biosynthetic gene clusters (BGCs)^9^. The BGCs of class I or class II bacteriocins typically include the precursor genes, and other associated genes such as transporter and peptidase (hereinafter referred to as context genes). Advanced by the rapid development of computational techniques, a range of dedicated bioinformatic tools have been in place for bacteriocin mining using distinct algorithms (Table 1). However, none of these tools are specifically designed for high-precision identification of class II bacteriocins. Even though class II bacteriocins belong to the group of antimicrobial peptides (AMPs) that do not undergo post-translational modifications, they need a unique mining approach compared to the general AMP prediction^25–27^. This is because while AMPs encompass entire small open reading frames, the maturation of most class II bacteriocins involves the removal of the leader peptide through peptidase, followed by secretion via transporter. Moreover, most tools predict class II bacteriocins solely by identifying precursor genes. Tools like RMSCNN^28^, BaPreS^29^, and BPAGS^30^ identify bacteriocin precursor sequences, but they do not differentiate between modified and unmodified bacteriocins. On the other hand, some tools, such as BAGEL4^31^ and PRISM 4^32^, concentrate only on precursor sequences similar to those of known bacteriocins. The most commonly used tool, antiSMASH^33^, has limited detection rules due to its focus on overall secondary metabolites, constraining a comprehensive identification of class II bacteriocins. Therefore, developing a more dedicated tool for class II bacteriocin mining is a prerequisite for interrogating them in the human gut microbiome.

**Table 1:**
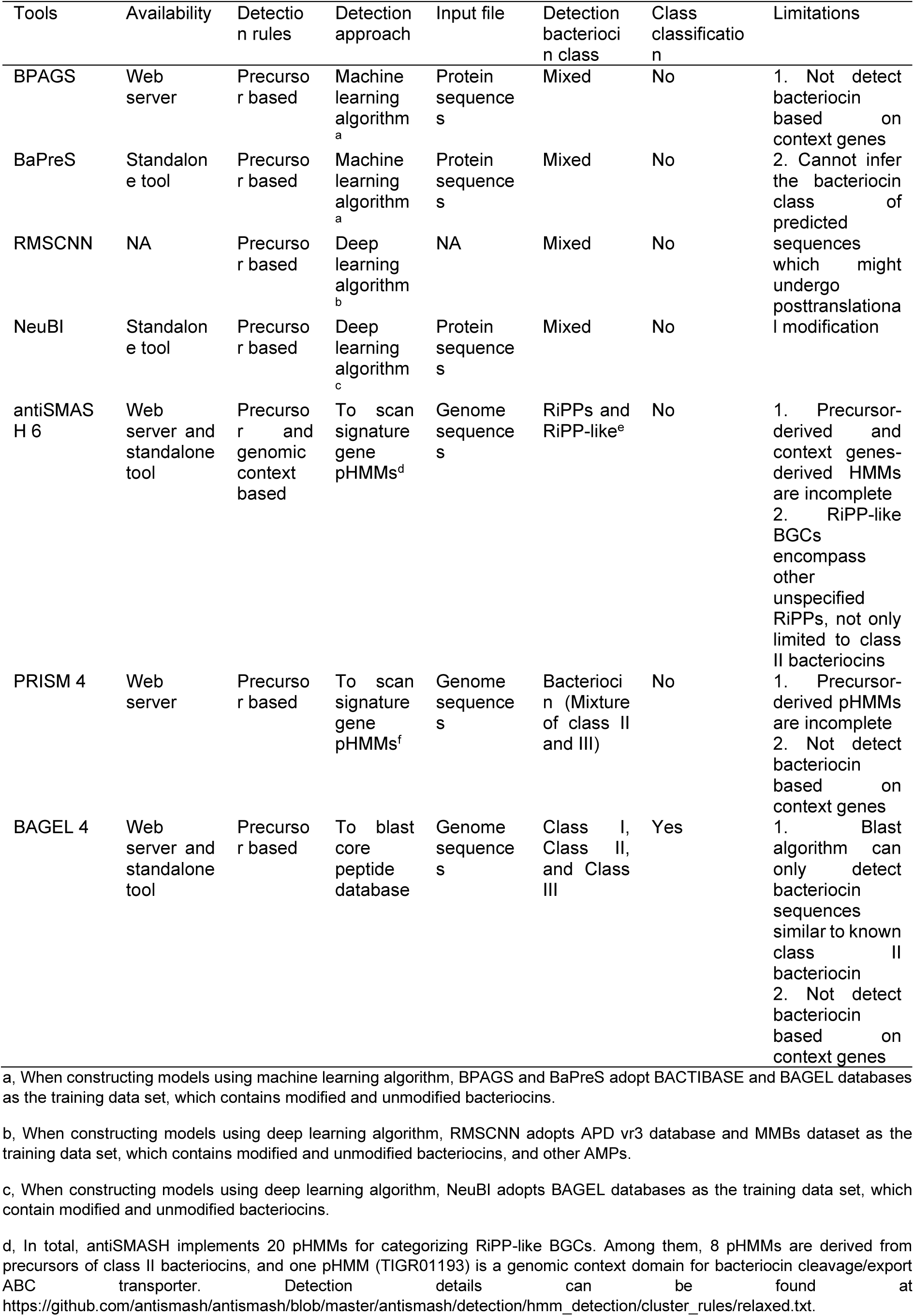
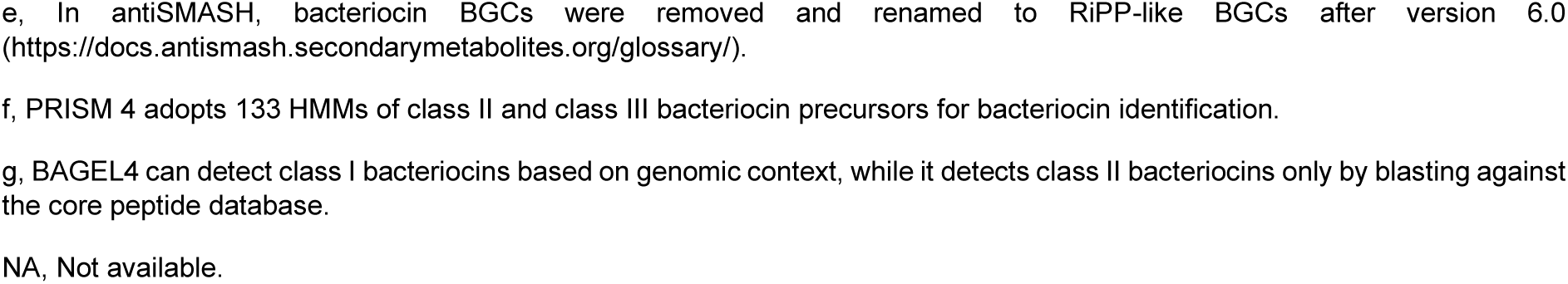
Comparison of current bioinformatic tools for bacteriocin identification.

| Tools | Availability | Detection rules | Detection approach | Input file | Detection bacteriocin class | Class classification | Limitations |
| --- | --- | --- | --- | --- | --- | --- | --- |
| BPAGS | Web server | Precursor based | Machine learning algorithm<sup>a</sup> | Protein sequences | Mixed | No | 1. Not detect bacteriocin based on context genes<br>2. Cannot infer the bacteriocin class of predicted sequences which might undergo posttranslational modification |
| BaPreS | Standalone tool | Precursor based | Machine learning algorithm<sup>a</sup> | Protein sequences | Mixed | No |  |
| RMSCNN | NA | Precursor based | Deep learning algorithm<sup>b</sup> | NA | Mixed | No |  |
| NeuBI | Standalone tool | Precursor based | Deep learning algorithm<sup>c</sup> | Protein sequences | Mixed | No |  |
| antiSMASH 6 | Web server and standalone tool | Precursor and genomic context based | To scan signature gene pHMMs<sup>d</sup> | Genome sequences | RiPPs and RiPP-like<sup>e</sup> | No | 1. Precursor-derived and context genes-derived HMMs are incomplete<br>2. RiPP-like BGCs encompass other unspecified RiPPs, not only limited to class II bacteriocins |
| PRISM 4 | Web server | Precursor based | To scan signature gene pHMMs<sup>f</sup> | Genome sequences | Bacteriocin (Mixture of class II and III) | No | 1. Precursor-derived pHMMs are incomplete<br>2. Not detect bacteriocin based on context genes |
| BAGEL 4 | Web server and standalone tool | Precursor based | To blast core peptide database | Genome sequences | Class I, Class II, and Class III | Yes | 1. Blast algorithm can only detect bacteriocin sequences similar to known class II bacteriocin<br>2. Not detect bacteriocin based on context genes |

a, When constructing models using machine learning algorithm, BPAGS and BaPreS adopt BACTIBASE and BAGEL databases as the training data set, which contains modified and unmodified bacteriocins.
b, When constructing models using deep learning algorithm, RMSCNN adopts APD vr3 database and MMBs dataset as the training data set, which contains modified and unmodified bacteriocins, and other AMPs.
c, When constructing models using deep learning algorithm, NeuBI adopts BAGEL databases as the training data set, which contain modified and unmodified bacteriocins.
d, In total, antiSMASH implements 20 pHMMs for categorizing RiPP-like BGCs. Among them, 8 pHMMs are derived from precursors of class II bacteriocins, and one pHMM (TIGR01193) is a genomic context domain for bacteriocin cleavage/export ABC transporter. Detection details can be found at https://github.com/antismash/antismash/blob/master/antismash/detection/hmm_detection/cluster_rules/relaxed.txt.
e, In antiSMASH, bacteriocin BGCs were removed and renamed to RiPP-like BGCs after version 6.0 (https://docs.antismash.secondarymetabolites.org/glossary/).
f, PRISM 4 adopts 133 HMMs of class II and class III bacteriocin precursors for bacteriocin identification.
g, BAGEL4 can detect class I bacteriocins based on genomic context, while it detects class II bacteriocins only by blasting against the core peptide database.
NA, Not available.

Here, we commenced with the development of a specialized tool called IIBacFinder (Class II Bacteriocin Finder), which integrates the detection of class II bacteriocins through both precursor genes and context genes. Demonstrated by recovering known class II bacteriocins from the genomes of bacteriocin producers, IIBacFinder outdoes the two wildly used tools, BAGEL4 and antiSMASH, in detecting class II bacteriocins. Alongside experimental validation, the systematic mining of ∼260,000 bacterial genomes from RefSeq database revealed the underappreciated biosynthetic potential of class II bacteriocins in the bacterial kingdom, particularly in Gram-negative bacteria. Those class II bacteriocins exhibited high niche-specificity and were enriched in the human microbiome. Leveraging the comprehensive analysis of ∼280,000 reference genomes reconstructed from human gut metagenomes, we gained deep insights into the biosynthetic capacity and chemical diversity of class II bacteriocins. Analyzing metagenomics and metatranscriptomics data of healthy human gut, we prioritized and chemically synthesized 26 predicted class II bacteriocins that were prevalent and abundant in healthy human gut microbiomes. They showed diverse narrow-spectrum antibacterial activities but had minimal impacts on human fecal microbiota in an *ex vivo* assay. Our work demonstrates the previously underappreciated biosynthetic potential of unmodified bacteriocins in the human gut microbiome, which might contribute to maintaining microbiome homeostasis and impacting human health.

## Results

### The development of IIBacFinder pipeline specializing in detecting unmodified bacteriocins

Several tools have been developed for bacteriocin identification with distinct mining strategies (Table 1), but they underperform in identifying class II bacteriocins primarily due to solely relying on precursor genes. To circumvent these limitations, we set out to develop an improved tool dedicated to identifying class II bacteriocins by integrating the detection of precursor genes and context genes. Given that profile hidden Markov models (pHMMs) offer a straightforward and rapid detection method to identify homologs within a specific protein family^34,35^, we constructed the pHMM library of precursor genes and context genes by curating known sequences and detecting their homologs from the UniRef100 database (Figure S1A, Tables S1, 2). Leveraging the library of 1,153 pHMMs of precursor genes and 37 pHMMs of context genes (Tables S3 - 5), we could comprehensively pinpoint class II bacteriocins by either directly detecting precursor genes or indirectly detecting novel bacteriocins genes through anchoring context genes (Figure 1A, Figure S1B). We termed this pipeline IIBacFinder (Class II Bacteriocin Finder).

**Figure 1.**
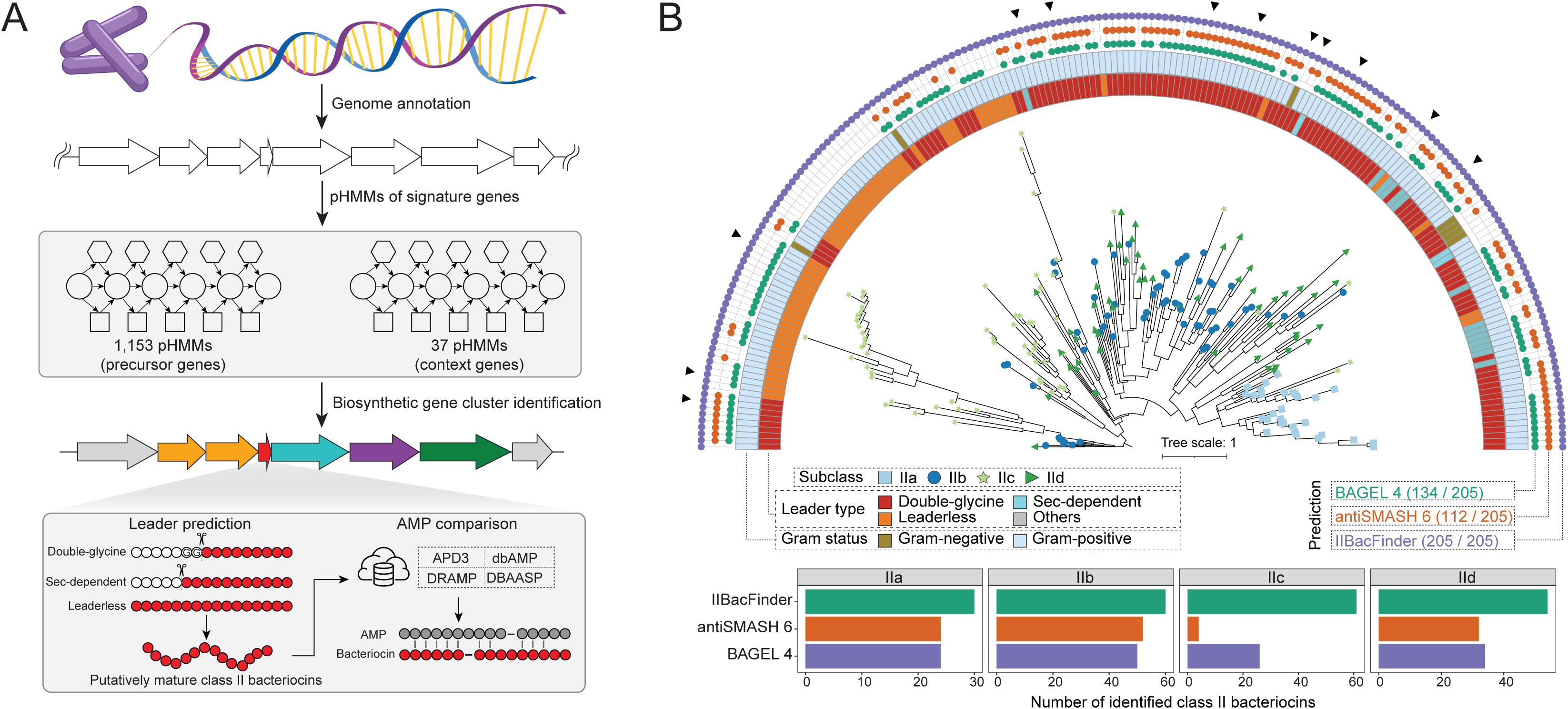
Overview of IIBacFinder specializing in detecting unmodified bacteriocins. (A) Schematic overview of the genome mining procedure in IIBacFinder. IIBacFinder detects potential unmodified class II bacteriocins from nucleotide sequences in FASTA format, by leveraging the library of 1,153 pHMMs of precursor genes and 37 pHMMs of context genes. The putative core peptides of hypothetical bacteriocins are deduced and compared against known AMPs from four databases (i.e., APD3, dbAMP v2.0, DRAMP v3.0, and DBAASP v3.0). (B) Performance of three tools in recovering 205 known class II bacteriocins from 137 bacterial genomes. The maximum likelihood phylogenetic tree was inferred based on full bacteriocin sequences. Symbols at the leaf tips represent the subclass of class II bacteriocins, and the black triangles indicate the 12 bacteriocin sequences which are not included when building pHMM models in IIBacFinder. Numbers in brackets indicate the number of bacteriocins detected by each prediction tool.

Following the detection of precursor genes, we sought to integrate the prediction of leader, which is a marked difference between class II bacteriocins and generic AMPs. Two types of leaders, the double-glycine leader and the sec-dependent leader, are generally present in class II bacteriocins. To streamline the prediction of double-glycine leader, we adopted and tuned a RiPP leader detection tool, NLPPrecursor^36^, for the cleavage site prediction. By employing 10-fold cross-validation, we determined that the modified algorithm could attain an accuracy of 98.0 % when factoring in cleavage points that are ±3 amino acids (AAs) away from the actual prediction site (Figure S1C, D). On the other hand, we applied SignalP 6.0^37^ to predict the cleavage site for the sec-dependent leader. Finally, the predicted core peptides were compared to known AMP sequences from four AMP databases^38–41^, to gain an insight into their sequence novelty (Figure 1A).

Upon the development of IIBacFinder, we next sought to evaluate its performance in identifying class II bacteriocins. We curated 137 bacterial genomes containing 205 complete precursor sequences of known class II bacteriocins, spanning different leader types and subclasses (Table S6). Among them, 191 sequences were unutilized when constructing the pHMM library of precursor genes, while 12 sequences were not because they were unintentionally excluded when building pHMMs. We then applied three tools (antiSMASH 6, BAGEL4, and IIBacFinder) to these genomes for detecting class II bacteriocins. Reassuringly, IIBacFinder could recover all bacteriocin precursor sequences (205/205, 100%), including 12 sequences that were new to IIBacFinder, while BAGEL4 and antiSMASH 6 detected 134 (65.4%) precursor sequences and 112 (54.6%) corresponding bacteriocin gene clusters, respectively (Figure 1B). Notably, their performance in identifying leaderless class IIc bacteriocins was significantly lower. The performance benchmarking demonstrated that IIBacFinder has a superior detection capability for unmodified class II bacteriocins compared to the other two commonly used tools.

### Underappreciated biosynthetic capacity of class II bacteriocins in the bacterial kingdom

Having attested to IIBacFinder’s remarkable ability to identify class II bacteriocins, we then applied it to 257,997 bacterial genomes from the NCBI RefSeq database^42^, in order to assess the global biosynthetic capacity of unmodified bacteriocins in the bacterial kingdom. These genomes spanned 56 phyla and 4,259 genera, corresponding to 96,325 Gram-positive and 161,472 Gram-negative bacteria. Using IIBacFinder, a total of 540,587 precursor sequences were identified from 133,806 genomes. Among them, 387,888 (71.75%) and 152,598 (28.23%) precursor sequences were detected from Gram-positive and Gram-negative bacteria, respectively. For comparison, antiSMASH 6 could detect 206,288 RiPP-like BGCs (encompassing BGCs encoding class II bacteriocins and other unclassified metabolites) in 119,697 genomes, consisting of 108,962 (52.84%) and 97,265 (47.16%) BGCs from Gram-positive and Gram-negative bacteria; meanwhile, BAGEL4 detected 137,942 precursor sequences with 93.36% being from Gram-positive genomes, indicating BAGEL4 biased the prediction in Gram-positive bacteria in which most known bacteriocins were discovered (Figure S2A, B).

To reduce the probability of false prediction, we only included 481,654 high-confidence sequences for downstream analysis (Figure S2C). The length of core peptides was approximately 40 amino acids for double-glycine and leaderless bacteriocins, but around 70 amino acids for sec-dependent bacteriocins (Figure S2D). Compared to known AMPs deposited in four databases (i.e., APD3, DRAMP, DBAASP, and dbAMP2), they showed less homology, with most being less than 40% identity (Figure S2E). Meanwhile, class II bacteriocins had higher median length than AMPs. By analyzing two physicochemical features, which play a pivotal role in elucidating the activity of AMPs, we found that double-glycine or leaderless bacteriocins exhibited a lower number of electronic charges than AMPs (Mann– Whitney U test, *P* < 0.001) but had higher GRAVY index scores, suggesting they are likely to be hydrophobic. In contrast, significantly lower GRAVY scores than AMPs implied that sec-dependent bacteriocins were more likely to be hydrophilic (Figure S2F).

We then conducted a rarefaction analysis on 8,610 precursor clusters that were categorized from 481,654 sequences using a 50% identity threshold, aiming to assess the genetically encoded biochemical diversity. The curve astonishedly showed that Gram-negative bacteria had a higher biosynthetic potential than Gram-positive bacteria (Figure 2A). We then decorated the GTDB bacterial tree with predicted precursor sequences to reveal the biosynthetic capacity at the phylum level. Through this examination, we found that the bacteriocin biosynthetic capacity stood out in several most sequenced phyla, such as Firmicutes (13,401 unique sequences), Proteobacteria (13,104), Bacteroidota (4,494), and Actinobacteriota (3,801) (Figure 2B). Among them, the bacteria from Firmicutes were the most prolific producers, with the average number of precursors per genome being 4.97, in line with the fact that most known bacteriocins have been discovered in this phylum. When down to the genus level, *Streptococcus*, as expected, topped the list of the bacterial tree of life. Other Gram-negative genera, such as the soil bacteria *Rhizobium* and *Bradyrhizobium*, also displayed promising biosynthetic capacity (Figure 2C). The overall survey indicated the previously underappreciated bacteriocin biosynthetic capacity in the bacterial kingdom, particularly in Gram-negative bacteria.

**Figure 2.**
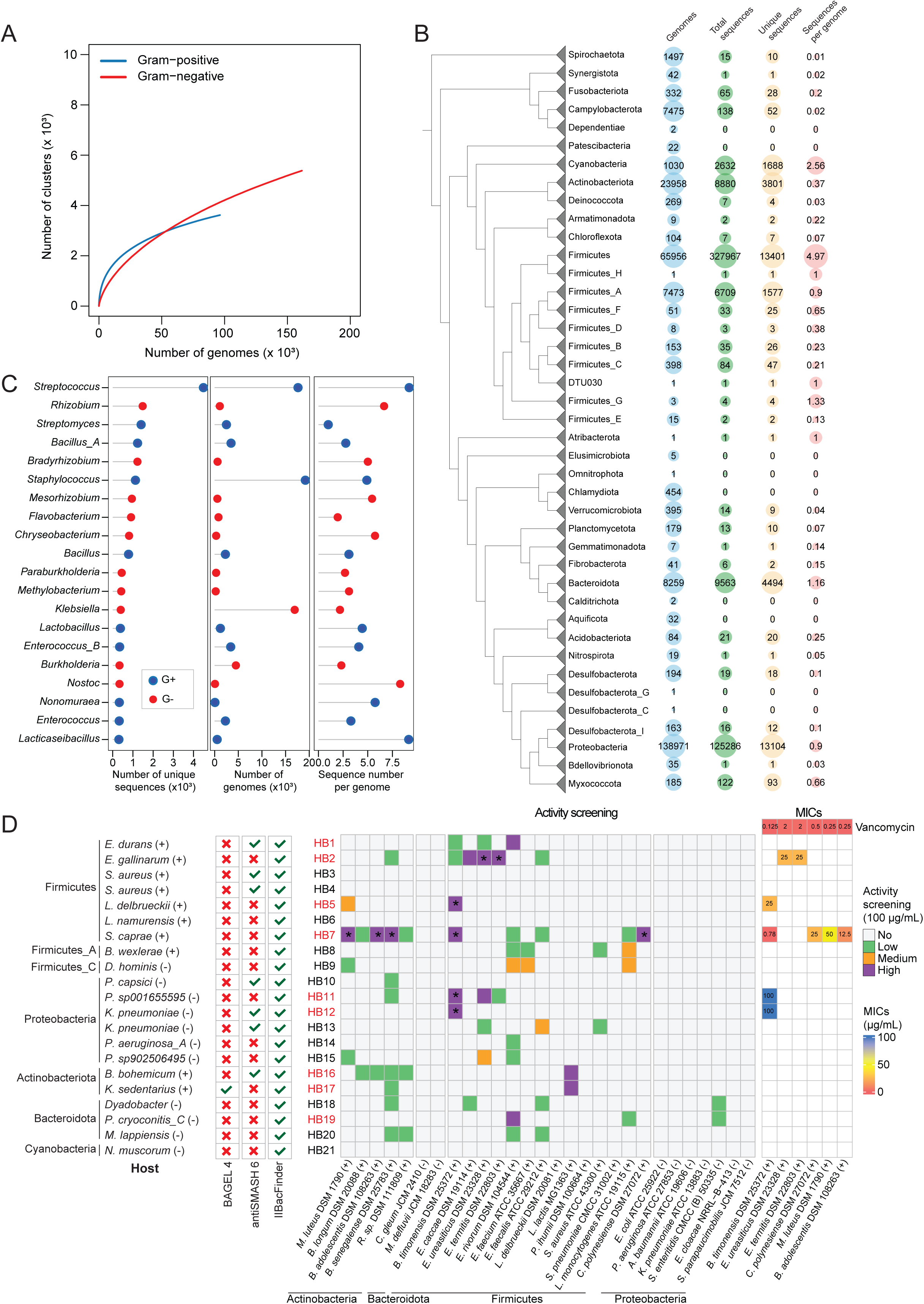
Biosynthetic potential of class II bacteriocins in the bacterial kingdom. (A) Rarefaction curves illustrating how the number of detected bacteriocins increases with the inclusion of more bacterial genomes. (B) The GTDB bacterial tree, decorated with the number of genomes, detected bacteriocin sequences, unique bacteriocin sequences, and bacteriocin sequences per genome, demonstrates the bacteriocin biosynthetic potential across different bacterial phyla. (C) The top 20 bacterial genera containing the most abundant class II bacteriocins. (D) Antibacterial activity of 21 synthetic hypothetical bacteriocins (HBs) against both Gram-positive (+) and Gram-negative (-) bacteria. A red cross signifies that HBs cannot be identified by the prediction tool, while a green tick indicates successful detection. The activity screening was conducted at a concentration of 100 µg/mL, with activity being categorized into four groups (no activity, < 25% inhibitory ratio or *P* value > 0.05; low activity, 25% ≤ inhibitory ratio < 50% & *P* value < 0.05; medium activity, 50% ≤ inhibitory ratio < 75% & *P* value < 0.05; high activity, inhibitory ratio > 75% & *P* value < 0.05). HBs with high activity are highlighted in red. Asterisks in the activity screening heatmap denote an inhibitory ratio of ≥ 90%, which is subject to further MIC examination (rightmost panel). Exact MIC values are indicated in the heatmap, with antibiotic vancomycin serving as the positive control. All assays are performed in three independent replicates.

To further demonstrate the prediction reliability, we arbitrarily selected and chemically synthesized 28 hypothetical bacteriocins (HBs) with the length of deduced core peptides < 50 amino acids for experimental validation, but 7 failed (Table S7). The remaining 21 sequences consisted of 13 double-glycine sequences, 5 sec-dependent sequences, and 3 leaderless sequences from seven phyla (i.e., Firmicutes, Firmicutes_A, Firmicutes_C, Cyanobacteria, Actinobacteria, Bacteroidota, Proteobacteria). Moreover, only HB17 could be detected by BAGEL4, while six HBs with double-glycine precursors could be captured by antiSMASH 6 as RiPP-like BGCs (Figure 2D). We examined their antibacterial potential against 19 Gram-positive and 9 Gram-negative indicator strains from four phyla at 100 µg/mL concentration (Figure 2D). The assay showed that 17 of 21 HBs could inhibit the growth of at least one indicator strain (inhibition ratio > 25%, Dunnett’s test *P* < 0.05). The active HBs primarily targeted the Gram-positive bacteria from phyla Firmicutes and Actinobacteria and barely inhibited Gram-negative bacteria. They displayed either narrow-spectrum activity (e.g., HB1 and HB2) or relatively broad-spectrum activity (e.g., HB7). Among them, 9 HBs (three from Gram-negative bacteria and six from Gram-positive bacteria) exhibited a robust activity with over 75% inhibition ratio, with minimal inhibitory concentrations (MICs) ranging from 0.78 to 100 µg/mL (Figure 2D). This suggests that IIBacFinder has the potential to identify genuine new unmodified bacteriocins, a capability that the other two tools lacked.

### Class II bacteriocins exhibit niche-specific enrichment in host-associated bacteria

The global inspection of class II bacteriocins revealed their wide distribution among diverse bacterial phyla, which might account for the previous concept that bacteriocin producers are widespread in natural environments^43^. To understand their biosynthetic potential in diverse ecosystems, we applied IIBacFinder to ∼50,000 metagenome-assembled genomes (MAGs) recovered from metagenomes of diverse habitats, including host-associated environments (primarily from human), aquatic environments, engineered environments, and terrestrial environments^44^. We identified 4,956 unique high-confidence precursor sequences from the genomic catalog of earth’s microbiomes, half of which (2,482 sequences) were from host-associated MAGs, followed by the engineered, aquatic, and terrestrial environments (Figure 3A). Notably, the bacteriocins identified from human contributed most to the host-associated bacteriocin (Figure 3B). Interestingly, we observed that class II bacteriocins exhibited high niche-specify, with >90 percent of unmodified bacteriocin sequences found in one specific ecosystem (Figure 3C).

**Figure 3.**
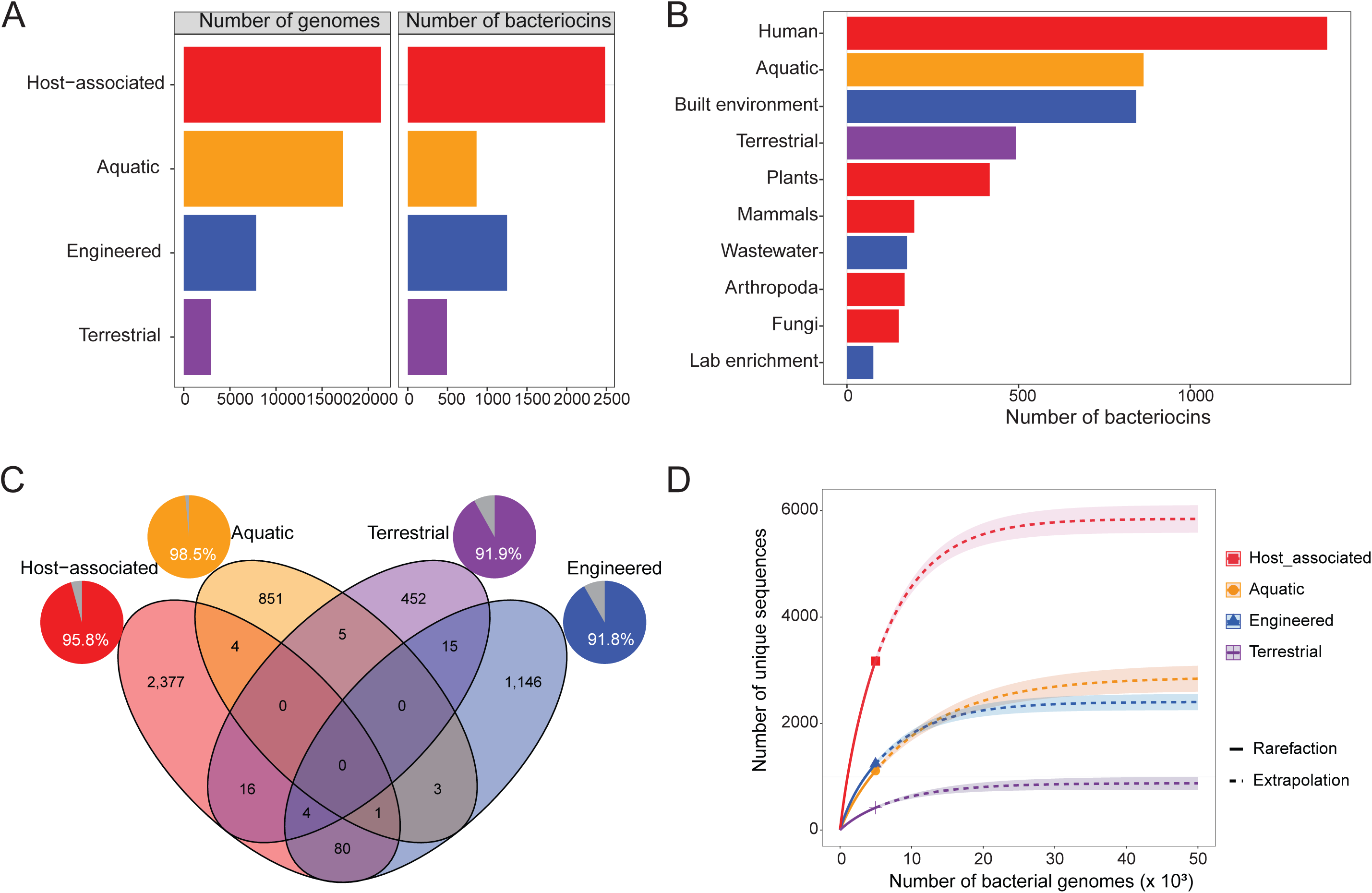
Class II bacteriocins exhibit niche-specific enrichment in host-associated bacteria. (A) Bar plots illustrating the number of genomes analyzed for class II bacteriocin identification and the corresponding number of non-redundant bacteriocin sequences detected. (B) Bar plot showing the number of non-redundant bacteriocin sequences detected from bacterial genomes across different habitats. (C) Venn diagram depicting the overlap of bacteriocin sequences detected from four distinct habitats, with pie charts indicating the proportion of niche-specific sequences, labeled with percentages. (D) Accumulation curves of bacteriocin sequences detected from the four habitats, with solid lines representing interpolated and actual data, and dashed lines representing extrapolated data.

Although the fact that the largest number of bacteriocins were detected in host-associated MAGs is informative, this might be skewed by the uneven number of reconstructed MAGs. Therefore, we then adopted rarefaction analyses to address this bias (Figure 3D). The rarefaction curve still indicated host-associated MAGs presented the greatest biosynthetic potential than other ecosystems, with being around three times over aquatic and engineered environments. The analysis of environmentally diverse bacteriocins implied class II bacteriocins’ potential to interact closely with community members in the host. They may also be utilized to interact with the host itself, particularly in the case of human hosts^20^.

### Human gut microbiota has the potential to produce diverse health-beneficial unmodified bacteriocins

Recognizing the prevalence of class II bacteriocins in the human microbiome, we focused on unmodified bacteriocin production in the human gut, where numerous microbes impact health by producing various metabolites^45^. We, therefore, applied IIBacFinder to ∼280,000 bacterial genomes recovered from the human gut microbiome^24^, resulting in 82,806 high-confidence precursor sequences, with 6,238 being unique (100% identity). More than half of them were double-glycine precursors (4,186, 67.10%), followed by sec-dependent leader precursors (765, 12.26%) and leaderless precursors (610, 9.78%). At the class level, Clostridia, Bacilli, and Bacteroidia were the most abundant producers. Despite not being predominant in the human gut microbiota, *Streptococcus* could produce the most diverse bacteriocins, aligning with its global bacterial biosynthetic potential. Other key human gut bacteria like *Prevotella*, *Faecalibacterium*, and *Bacteroides* also exhibited high biosynthetic potential (Figure 4A).

**Figure 4.**
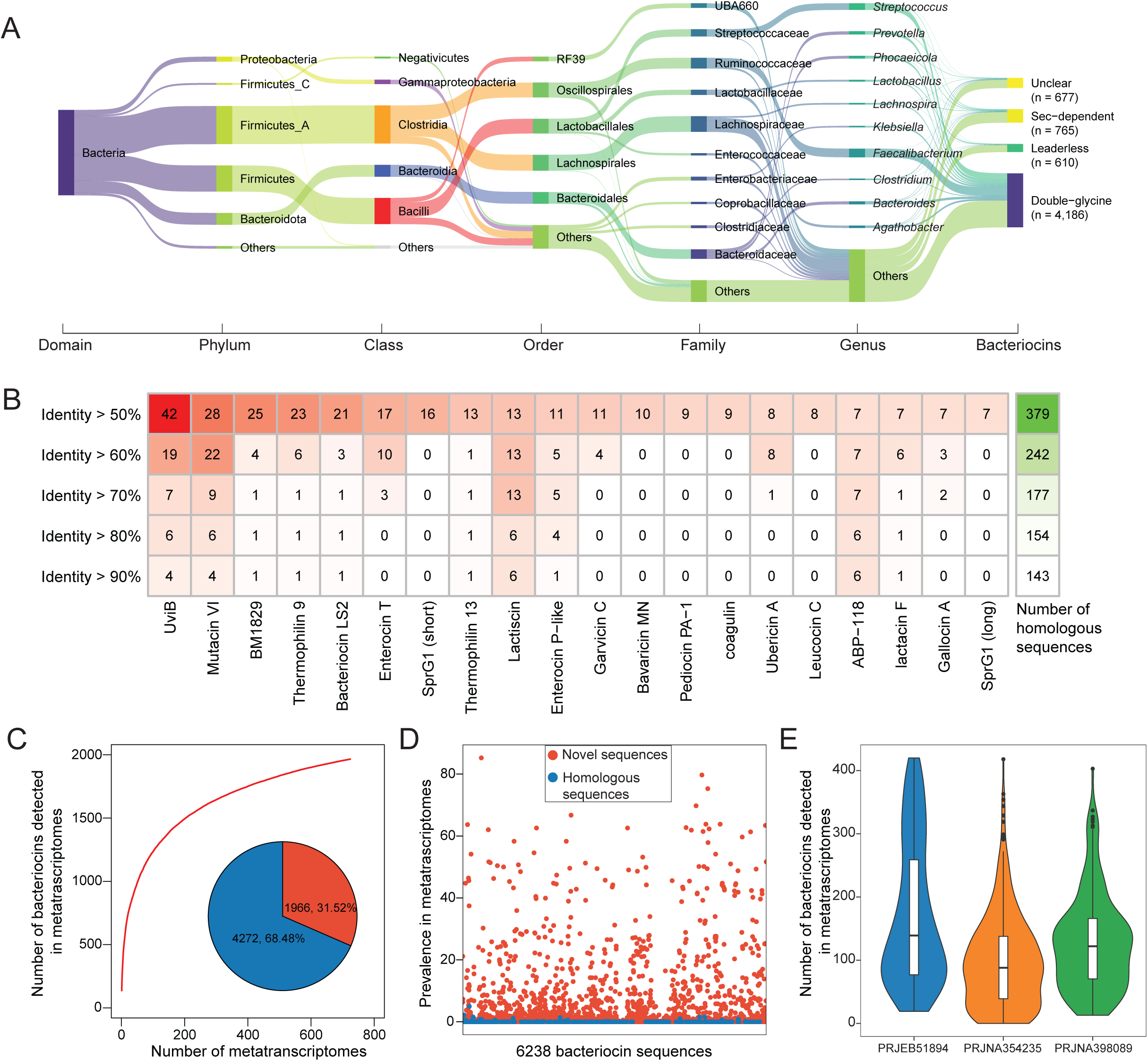
Human gut bacteria have significant potential to produce a diverse array of unmodified bacteriocins. (A) Sankey plot showing the bacteriocin biosynthetic potential of 285,835 human gut bacteria across different taxonomic levels. Numbers in brackets indicate the number of non-redundant class II bacteriocin sequences detected. (B) Heatmap illustrating the number of gut microbiota-derived class II bacteriocins homologous to specific (left panel) and all (right panel) known class II bacteriocins at varying identity thresholds (50%-90%). Only the top 20 most abundant known class II bacteriocins are shown. (C) Rarefaction curves showing the increase in detected bacteriocins with the inclusion of metatranscriptomes. The pie chart indicates the percentage of class II bacteriocins detected (red) and not detected (grey) in 724 metatranscriptomes. (D) Scatter plot illustrating the prevalence of 6,238 putative bacteriocins in metatranscriptomes. (E) Violin plot depicting the distribution of the number of class II bacteriocins actively transcribed in individuals from three cohorts.

Although this analysis indicated the high biosynthetic potential of human gut bacteria, only a few class II bacteriocins have been identified from gut microbiota yet^18^. We, thus, wondered whether these predicted unmodified bacteriocins were chemically distinct from known ones. We thus compared the 6,238 unique sequences with 298 known bacteriocin sequences at varying identity thresholds of 50% - 90% (Figure 4B, Figure S3). To our surprise, we found 379 (6.07%) and 143 (2.29%) homologous sequences at the identity of 50% and 90%, respectively, indicating that gut microbiota could produce similar bacteriocin products, but the majority remained uncharacterized. Among the most diverse were UviB homologies produced by *Clostridium perfringen*^46^. Some bacteriocins have established functions in mediating host health, for example, a broad-spectrum class II bacteriocin, ABP118, can protect mice against infection with pathogen *L. monocytogenes*^47^. These findings suggest that unmodified bacteriocins with protective functions may have a wide range of health-promoting effects.

Recent advancements in functional metagenomics have facilitated the discovery of bioactive RiPPs in the largely uncharted human microbiota, and have established links to their potential physiological roles^48^. We next looked at evidence of their transcription in 724 metatranscriptomes of healthy individuals from three cohorts to look for insight into their possible physiological roles. We found 1,966 (31.52%) sequences were actively transcribed in at least one sample, but more sequences could be detected with an increase in sample size as indicated by rarefaction analysis (Figure 4C). Most of actively transcribed bacteriocins are predicted novel (Figure 4D), might involving uncharted roles in the human gut. Furthermore, multiple distinct bacteriocin sequences were found to be transcribed in each individual, with a median number of 139, 88, and 122 in three cohorts (Figure 4E), implying that hundreds of transcribed bacteriocins might create a protective barrier for pathogen invasion in a synergistic manner^49^.

### Omics analysis-guided prioritization and identification of unmodified bacteriocins from the healthy gut microbiome

Inspecting the class II bacteriocins from human gut MAGs allowed us to appreciate their significant biosynthetic potential in the human gut. Next, we set out to experimentally characterize the bioactive class II bacteriocins that might play a crucial role in maintaining healthy human gut homeostasis. As the species abundance and sequence depth would skew the genome assembly from metagenomics data, we prioritized putatively bioactive bacteriocin using mapping-based analysis of metagenomic and metatranscriptomic data instead of those that were the most abundant in gut MAGs. To do so, we first created a reference database of class II bacteriocin by applying IIBacFinder to ∼700,000 bacterial genomes from multiple sources, including RefSeq^42^, human bodies^24,50,51^, earth environment^44^, ocean^52^, mouse and ruminant gastrointestinal tract^53,54^. Following the construction of this database encompassing 32,735 non-redundant high-confidence class II bacteriocin sequences, we examined their profiles by re-visiting 1,901 metagenomic and 724 metatranscriptomic datasets from healthy individuals (Figure 5A). To remove the redundancy, we computed their abundance at the cluster level (80% identity and 80% coverage) rather than at the individual level, and 3,581 bacteriocin clusters were detected in this omics analysis. Given the fact that bioactive bacteriocin could inhibit the growth of other bacteria and thus might regulate the overall bacterial community, we ranked them by considering (1) their prevalence and (2) abundance, (3) negative association with bacterial species, and (4) association with bacterial community diversity assessed by Shannon index (Figure 5A). This analysis allowed us to prioritize bioactive bacteriocins in the human gut microbiome (Table S8).

**Figure 5.**
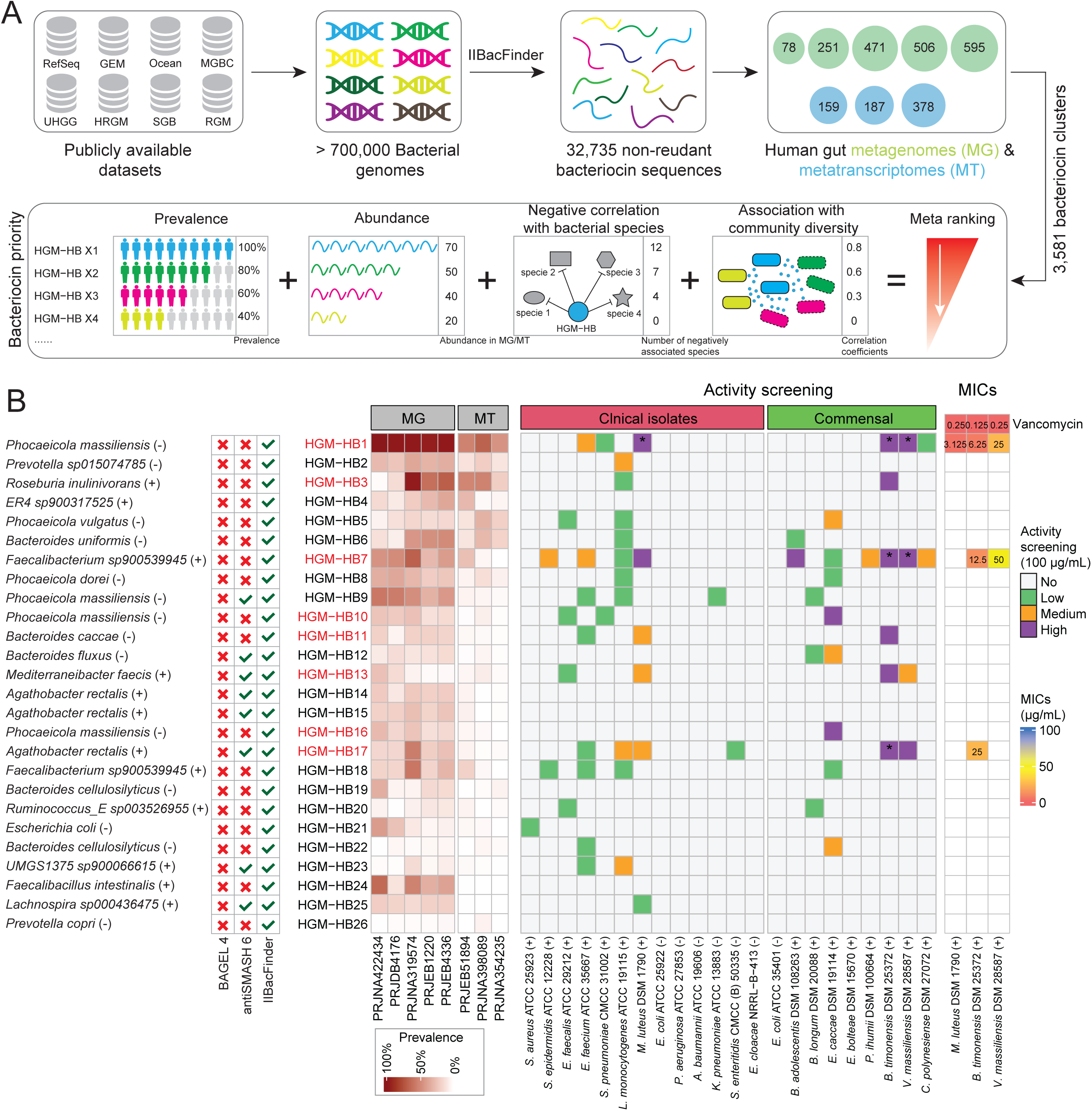
Prioritization and experimental validation of class II bacteriocins from human gut microbiota. (A) Schematic overview of prioritizing class II bacteriocins using metagenomics and metatranscriptomics analysis. Class II bacteriocins are detected from >700,000 bacterial genomes, which are subsequently surveyed in human gut metagenomes (MG) and metatranscriptomes (MT). A total of 3,581 bacteriocin clusters detected gut MG/MTare ranked based on a meta-ranking scheme consisting of four criteria (i.e., prevalence, abundance, negative association with bacterial species, and correlation with community diversity). (B) Antibacterial activity of 26 human gut microbiota-derived hypothetical bacteriocins (HGM-HBs). A red cross signifies that HGM-HBs cannot be identified by the prediction tool, while a green tick indicates successful detection. The heatmap in the middle panel depicts the prevalence and abundance of class II bacteriocins in different cohorts. The ones in the right panel show the activity screening for 26 HGM-HBs at a concentration of 100 µg/mL. HGM-HBs with high activity are highlighted in red. Asterisks in the activity screening heatmap denote an inhibitory ratio of ≥ 90%, which is subject to further MIC examination (rightmost panel). Exact MIC values are indicated in the heatmap, with antibiotic vancomycin serving as the positive control. All assays are performed in three independent replicates.

Finally, on the basis of the ranking analysis, we scrutinized and selected 40 clusters of human gut microbiota-derived hypothetical bacteriocins (HGM-HBs) with a length of core peptide < 50 amino acids for experimental validation (see STAR Methods). We attempted to chemically synthesize the core peptides from their representative sequences; however, 14 of these attempts were unsuccessful. Consequently, we had 26 HGM-HBs available for examination (Table S9). All of them were identified from human gut-derived bacterial genomes. None was detected by BAGEL 4, while eight were identified by antiSMASH 6 (seven within RiPP-like BGCs and one within NRPS BGC). Meanwhile, HGM-HP11 was identical to bacteroidetocin A^55^, but we did not include the intrinsic disulfide bonds for chemical synthesis as the putative disulfide bonds in other HGM-HPs were not predictable. Likewise, we initially examined their antibacterial activity against 21 indicator strains encompassing 13 clinical pathogens and 8 gut commensal bacteria at a concentration of 100 µg/mL. Under a cutoff of inhibition ratio > 25% and *P* < 0.05, 16 peptides (70%) were found to be active against, generally, Gram-positive bacteria. Notably, 8 HGM-HBs, including HGM-HP11, effectively inhibited at least one indicator strain with a > 75% inhibition ratio. Further activity examination for three HGM-HBs (HGM-HP1, HGM-HP7, and HGM-HP17) showed their MICs ranging from 3.125 to 50 µg/mL (Figure 5B). The *in vitro* assay indicated that the human gut microbiota-derived class II bacteriocin could inhibit the growth of clinical pathogens and gut commensals.

### Narrow-spectrum bacteriocins tweak human fecal microbiota

Although *in vitro* activity screening against a limited panel of indicator strains demonstrated active unmodified bacteriocins from the human gut microbiome, whether and how they affect the entire gut microbial community remains unclear. To address this, we utilized a human fecal-derived *ex vivo* microbial community to model the effects of bacteriocins on gut microbiota (Figure 6A). We tested three active bacteriocins (HGM-HP1, HGM-HP7, and HGM-HP17) and an inactive one (HGM-HP14) as determined by *in vitro* assay (Figure 5B). Vancomycin was included as a positive control. We investigated the effects of two concentrations, 1 μg/mL and 100 μg/mL, to understand how these bacteriocins modulate the microbial community at a low concentration, and to determine if they exhibit a broader activity spectrum than previously observed *in vitro* which could be indicated at a high concentration. Following a 48-hour treatment with bacteriocin peptides or vancomycin, no pronounced change in fecal microbiota composition was observed, except with 100 μg/mL of vancomycin (Figure 6B). Additionally, only the 100 μg/mL concentration of vancomycin, and not the other treatments, significantly decreased microbial diversity as indicated by the Shannon diversity index (Figure 6C), and markedly altered the microbial community structure as shown by principal coordinate analysis based on Bray-Curtis distances; however, 100 μg/mL of HGM-HP14 and HGM-HP17 slightly changed the overall microbial community (Figure 6D, Figure S4A, B). These results suggest that putative bacteriocins have only slight effects on human fecal microbiota, even at high concentrations.

**Figure 6.**
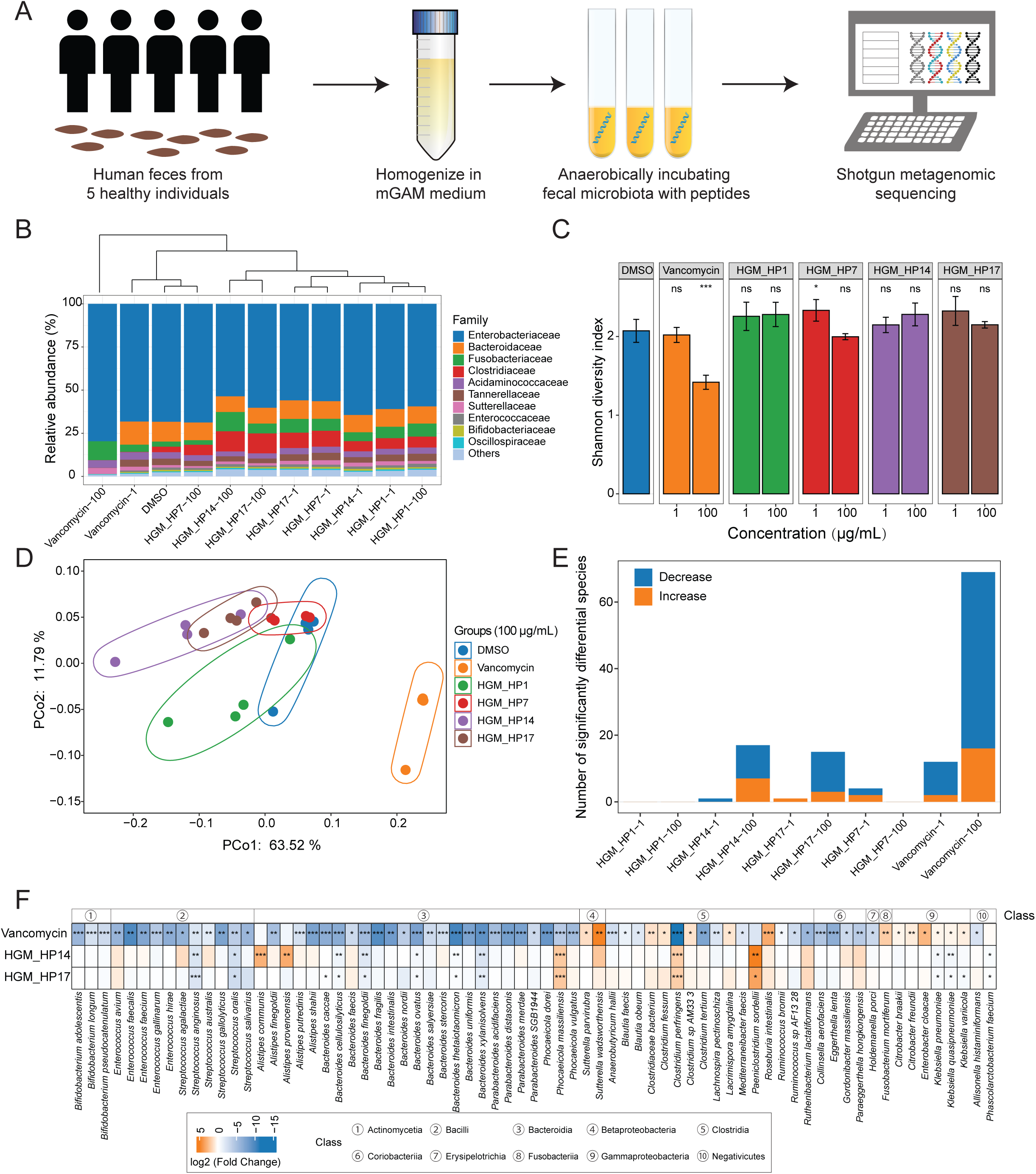
Narrow-spectrum bacteriocins tweak human fecal microbiota in *ex vivo* assay. (A) Schematic overview of the *ex vivo* assay to explore the impact of bacteriocins on human fecal microbiota. (B) Stacked bar plot displaying the average relative abundance of bacterial families, with a top dendrogram illustrating hierarchical clustering based on the Bray–Curtis dissimilarity of microbial communities at the family level. (C) Bar plot depicting the alpha diversity of the microbial community at the species level, measured by the Shannon diversity index. Data are presented as mean ± standard deviation. Statistical significance between treatment groups and the control group (DMSO) was determined using a two-sided Welch’s *t*-test. (D) Principal Coordinate Analysis (PCoA) of the microbial community with the treatment of 100 µg/mL of vancomycin or bacteriocins, based on Bray-Curtis dissimilarity at the species level. (E) Stacked bar plot showing the number of significantly differential species identified by MaAsLin2. (F) Heatmap representing the 71 significantly differentially abundant species between three treatment groups (100 µg/mL of vancomycin, HGM_HP14, and HGM_HP17) vs. the DMSO group. Each treatment group is symbolized by the bacteriocin and the concentration tested (e.g., HGM_HP1-1 represents 1 µg/mL of HGM_HP1, whereas HGM_HP1-100 represents 100 µg/mL of HGM_HP1). Statistical significance is denoted as follows: \**P* < 0.05; \*\**P* < 0.01; \*\*\**P* < 0.001; ns, not significant.

We then used MaAsLin2^56^ to examine the abundance difference in bacterial species between treatment and control groups, identifying a variable number of species that could be affected by bacteriocins or vancomycin (FDR-adjusted *P* value < 0.05) (Figure 6E, Figure S4C). Results indicated that a low concentration (1 μg/mL) of bacteriocins had minimal impact on fecal bacterial species. However, at a high concentration (100 μg/mL), two bacteriocins, HGM-HP14 and HGM-HP17, significantly affected the abundance of 17 and 15 bacterial species, respectively (Figure 6E). Notably, HGM-HP14, which showed no activity *in vitro*, decreased the abundance of 10 bacterial species in the larger microbial community, suggesting that *in vitro* screening might overlook some authentic bacteriocins. In contrast, the other two active bacteriocins (HGM-HP1 and HGM-HP7) had negligible effects on fecal microbiota, even at 100 μg/mL. As expected, the broad-spectrum antibiotic vancomycin significantly affected numerous bacterial species, particularly at the 100 μg/mL concentration, where it decreased the abundance of 53 species (Figures 6E, F). Vancomycin primarily reduced species from the classes Bacilli and Bacteroidia. Similarly, the narrow-spectrum bacteriocins HGM-HP14 and HGM-HP17 also reduced several bacterial species from these classes. Taken together, the *ex vivo* assay demonstrated slight effects of narrow-spectrum bacteriocins on human fecal microbiota.

## Discussion

The human gut microbiota is a rich source of bacteriocins that might combat pathogenic bacteria invasion and participate in human physiology^18,19^. Although a few efforts have been made to decipher their distribution in human gut microbiome^22,23^, the unmodified bacteriocins remain incompletely characterized. In this study, we conducted the first large-scale survey, to our best knowledge, on unmodified class II bacteriocins in the bacterial kingdom and human gut microbiome by applying an in-house bioinformatic tool. Through systematic investigation in combination with experimental validation, we demonstrated novel class II bacteriocins throughout the tree of bacterial life as well as in the human gut microbiome. These investigations will not only disclose that the class II bacteriocins are largely untapped resources for antimicrobial discovery but also act as a crucial pathway for unraveling bacteriocins-mediated interactions between the microbiome and human host.

Antimicrobial resistance (AMR) is a major global health concern that requires immediate, effective responses. Bacteriocins, due to their diversity, high efficacy, and ease of bioengineering, are emerging as potential alternatives to traditional antibiotics^7^. They often specifically target pathogens, thus limiting the damage caused by broad-spectrum antibiotics to existing microbial communities and potentially reducing adverse health effects^57^. For instance, Thuricin CD targets *Clostridium difficile* with minimal impact on the microbial community, yet with efficacy comparable to broad-spectrum antibiotics^58^. Our study found that class II bacteriocins exhibit narrow-spectrum activity, suggesting their potential to target pathogens specifically. Unmodified class II bacteriocins are preferred over modified counterparts for heterologous production and chemical synthesis, making them more practical for large-scale antimicrobial discovery efforts. In this regard, big data omics mining offers a faster and more high-throughput approach than traditional screening methods. To enhance the detection of class II bacteriocins, we developed a specialized software pipeline that identifies them by scanning pHMMs of signature genes. This advanced tool has unveiled the untapped biosynthetic potential for class II bacteriocins within the bacterial kingdom and the human gut microbiome, facilitating their large-scale discovery. Their genomics-guided discovery could be achieved by cell-free protein synthesis systems or robotic automation systems; two techniques have been successfully applied to AMP identification and modified bacteriocin characterization^59–61^. Hence, we believe that future advances in techniques can accelerate the discovery of class II bacteriocins, providing a promising strategy to address the AMR issue.

Recent studies have uncovered and experimentally confirmed a multitude of AMPs originating from the global microbiome^25^ and the human gut microbiome^26,27^, highlighting the overlooked potential of antagonistic AMPs within the bacterial realm, particularly within the human gut microbiota. While there are still questions about whether these AMPs are genuinely produced, secreted, and undergo modifications similar to bacteriocin biosynthesis, they hold potential as new bacteriocins. Unlike previous studies, our research primarily focuses on unmodified bacteriocins that are likely naturally produced and secreted. We pay particular attention to mature peptides, as opposed to the full coding sequences typically examined in AMP prediction. We have determined that synthetic bacteriocins, originating from the human gut microbiome, demonstrate antibacterial properties. However, their *in vivo* functions might extend beyond this, as they also display additional antimicrobial activities such as combating biofilms, fungi, and viruses^2,64^. For instance, enterocin A inhibits biofilm formation in three known pathogens, which can aid in reducing harmful pathogens in the human gut^58^. Furthermore, plantaricins EF and JK^11^, known for their antifungal properties, could influence human health indirectly by regulating the vital fungal community^62^. On top of antimicrobial activities, unmodified bacteriocins can also directly interplay with the human host, exhibiting various activities such as anticancer^58^ and immunomodulation^63–65^. Notably, plantaricin EF increases the production of proteins that fortify the intestinal lining, preserving its integrity^15^. Our comprehensive review reveals that human gut bacteria can produce a variety of class II bacteriocins, suggesting their potential roles in promoting human health. Consequently, we advocate for considering the role of class II bacteriocins in studies exploring the relationship between gut microbiota and human health.

In conclusion, we have developed an advanced tool, IIBacFinder, to provide a comprehensive overview of unmodified bacteriocins within the bacterial kingdom, with a particular focus on the human gut microbiome. Our study reveals the previously underestimated chemical repertoire of unmodified bacteriocins, revealing an untapped treasure for antimicrobial discovery. Diverse class II bacteriocins encoded by the human gut microbiome imply their under-perceived roles that cannot be ignored in future studies. Thus, our study paves the way for antimicrobial discovery from the untapped human microbiome and for deepening our understanding of its role in human health and disease in future studies.

## Limitations of the study

Identifying bacteriocins produced by human microbiota is crucial for comprehending the communication mechanisms within gut microbiota and between the microbiota and the human host. Although our tool, IIBacFinder, significantly improves the identification of unmodified class II bacteriocins, it may still yield false predictions, especially when detecting new bacteriocins based on context genes. Due to the limited number of characterized class II bacteriocins, our current understanding makes it difficult to differentiate real bacteriocins from false predictions without experimental validation. Moreover, while we can identify new bacteriocins through anchoring context genes, we may miss other unknowns that don’t fit our current understanding of class II bacteriocin biosynthetic logic, leading to an incomplete exploration of their biosynthetic potential. Future advancements in high-confidence techniques are needed to identify authentic bacteriocins accurately.

In this study, we chemically synthesized predicted mature peptides of unmodified bacteriocins, which may not accurately represent their natural mature form. Despite not undergoing post-translational modifications, unmodified bacteriocins might have other intramolecular features, like disulfide bonds and a reduced state of cysteine^10,66^. Additionally, bacteriocin synergies might be common in the human gut^49^, considering that hundreds of bacteriocins are actively transcribed in individuals and many bacteriocin gene clusters often contain more than one precursor gene. Thus, the *in vivo* functions of synthetic bacteriocins might differ from our *in vitro* characterization. Other potential functions of these bacteriocins remain unknown and require further investigation. Despite these limitations, our study emphasizes that the gut microbiota could produce a variety of class II bacteriocins that might interact with the microbial community or human host.

## Resource availability

### Lead contact

Requests for further information and resources should be directed to and will be fulfilled by the lead contact, Yong-Xin Li.

### Materials availability

This study did not generate new unique reagents.

### Data and code availability

- The bacterial genomes were collected from eight publicly available datasets, including the NCBI Assembly RefSeq database (https://ftp.ncbi.nlm.nih.gov/genomes/refseq/), human microbiome (http://segatalab.cibio.unitn.it/data/Pasolli_et_al.html), UHGG database (, https://www.ebi.ac.uk/ena/browser/view/PRJEB33885), HRGM database (https://www.mbiomenet.org/HRGM/), Earth’s microbiome (https://portal.nersc.gov/GEM/genomes/), ocean microbiome (https://www.ebi.ac.uk/ena/browser/view/PRJEB45951), mouse (https://zenodo.org/records/4840600) and ruminant gut microbiome (https://www.ebi.ac.uk/ena/browser/view/PRJNA657473).
- The raw data for metagenomes and metatranscriptomes are publicly available in NCBI-SRA under the BioProject IDs: PRJNA422434, PRJNA319574, PRJEB4336, PRJEB1220, PRJDB4176, PRJNA354235, PRJEB51894, and PRJNA398089.
- Publicly available AMP data can be accessed from four databases – APD3: https://aps.unmc.edu/downloads; dbAMP v2.0: https://awi.cuhk.edu.cn/dbAMP/download.php; DRAMP v3.0: http://dramp.cpu-bioinfor.org/downloads/; DBAASP v3.0: https://dbaasp.org/.
- Protein family models can be found in the Pfam database (version 34.0, http://ftp.ebi.ac.uk/pub/databases/Pfam/releases/Pfam34.0) and NCBI database (version 11.0, https://ftp.ncbi.nlm.nih.gov/hmm/11.0/).
- Other source data required to reanalyze the data reported in this paper can be found in supplementary materials or the Zenodo repository (https://zenodo.org/records/13588376).
- Raw read sequences of the shotgun metagenomic sequences generated in this study were deposited at Sequence Read Archive (SRA, NCBI) under the accession of PRJNA1201247.
- The IIBacFinder codes can be found at https://github.com/ZhangDengwei/IIBacFinder and https://zenodo.org/records/14292149.

## Acknowledgments

The author would like to thank Prof. Wang (University of Nebraska Medical Center) for graciously providing the AMP sequences from the latest APD3 database in response to an email request. The authors also thank Vecteezy.com for distributing free vector art at https://www.vecteezy.com, which is partially adopted and modified in the graphical abstract.

## Author contributions

Conceptualization: Y.L. and D.Z. Methodology: D.Z., Y.Z., Y.S., J.Z., J.L., G.W., J.Z., Y.G. Investigation: Y.L. and M.C. Visualization: Y.L. and D.Z. Supervision: Y.L. and M.C. Writing— original draft: D.Z. Writing—review & editing: Y.L. and D.Z.

## STAR Methods

### Key resources table

| REAGENT or RESOURCE | SOURCE | IDENTIFIER |
| --- | --- | --- |
| **Bacterial and virus strains** |  |  |
| *Bacillus timonensis* | Leibniz Institute DSMZ | DSM 25372 |
| *Enterococcus caccae* | Leibniz Institute DSMZ | DSM 19114 |
| *Enterococcus ureasiticus* | Leibniz Institute DSMZ | DSM 23328 |
| *Enterococcus termitis* | Leibniz Institute DSMZ | DSM 22803 |
| *Enterococcus rivorum* | Leibniz Institute DSMZ | DSM 104544 |
| *Enterococcus faecium* | American Type Culture Collection | ATCC 35667 |
| *Enterococcus faecalis* | American Type Culture Collection | ATCC 29212 |
| *Lactobacillus delbrueckii* | Leibniz Institute DSMZ | DSM 20081 |
| *Lactococcus lactis* subsp. *cremoris* | N/A | MG1363 |
| *Paenibacillus ihumii* | Leibniz Institute DSMZ | DSM 100664 |
| *Staphylococcus aureus* | American Type Culture Collection | ATCC 43300 |
| *Streptococcus pneumoniae* | N/A | CMCC 31002 |
| *Listeria monocytogenes* | American Type Culture Collection | ATCC 19115 |
| *Clostridium polynesiense* | Leibniz Institute DSMZ | DSM 27072 |
| *Staphylococcus aureus* | American Type Culture Collection | ATCC 25923 |
| *Staphylococcus epidermidis* | American Type Culture Collection | ATCC 12228 |
| *Virgibacillus massiliensis* | Leibniz Institute DSMZ | DSM 28587 |
| *Enterocloster bolteae* | Leibniz Institute DSMZ | DSM 15670 |
| *Escherichia coli* | American Type Culture Collection | ATCC 25922 |
| *Escherichia coli* | American Type Culture Collection | ATCC 35401 |
| *Pseudomonas aeruginosa* | American Type Culture Collection | ATCC 27853 |
| *Acinetobacter baumannii* | American Type Culture Collection | ATCC 19606 |
| *Klebsiella pneumoniae* | American Type Culture Collection | ATCC 13883 |
| *Salmonella enteritidis* | N/A | CMCC (B) 50335 |
| *Enterobacter cloacae* | N/A | NRRL-B-413 |
| *Sphingomonas parapaucimobilis* | Japan Collection of Microorganisms | JCM 7512 |
| *Micrococcus luteus* | Leibniz Institute DSMZ | DSM 1790 |
| *Bifidobacterium longum* subsp. *infantis* | Leibniz Institute DSMZ | DSM 20088 |
| *Bifidobacterium adolescentis* | Leibniz Institute DSMZ | DSM 108263 |
| *Brevibacterium senegalense* | Leibniz Institute DSMZ | DSM 25783 |
| *Rothia* sp. | Leibniz Institute DSMZ | DSM 111809 |
| *Chryseobacterium gleum* | Japan Collection of Microorganisms | JCM 2410 |
| *Mucilaginibacter defluvii* | Japan Collection of Microorganisms | JCM 18283 |
| **Chemicals, peptides, and recombinant proteins** |  |  |
| Brain heart infusion medium (BHI) | hopebio | HB8297-1 |
| Tryptic Soy Broth (TSB) | hopebio | HB4114 |
| De Man–Rogosa–Sharpe medium (MRS) | BEIJING LAND BRIDGE | CM187 |
| Peptone Yeast Glucose medium (modified) (mPYG) | TUOPU BIOTECHNOLOGY | MD090 |
| Reinforced Clostridium Medium (RCM) | hopebio | HB0316 |
| Gifu Anaerobic Broth (GAM) | TUOPU BIOTECHNOLOGY | M5650 |
| Bifidobacterium Medium (BM) | hopebio | HB8527 |
| M17 broth | Solarbio | LA0610 |
| Deposited data |  |  |
| Code for IIBacFinder | This study | https://github.com/ZhangDengwei/IIBacFinder |
| Metagenomic sequencing data of human feces | This study | PRJNA1201247 |
| Software and algorithms |  |  |
| Python 3.7.12 | Python Software Foundation | https://www.python.org/ |
| R version 4.2.2 | The R Foundation | https://www.r-project.org/ |
| Prodigal-short | Santos-Aberturas et al.<sup>67</sup> | https://github.com/streptomyces/ripper/tree/master |
| CD-HIT 4.8.1 | Fu et al.<sup>68</sup> | https://github.com/weizhongli/cdhit |
| numpy |  |  |
| pandas |  |  |
| HMMer 3.1b2 | Sean R. Eddy<sup>69</sup> | http://hmmer.org/ |
| Ampir |  |  |
| amPEPpy 1.0 |  |  |
| SAMtools 1.18 |  |  |
| Diamond v2.1.8.162 | Buchfink et al.<sup>70</sup> | https://github.com/bbuchfink/diamond |
| MAFFT v7.508 | Katoh et al.<sup>71</sup> | https://mafft.cbrc.jp/alignment/software/ |
| trimAl v1.4.rev15 | Capella-Gutiérrez et al.<sup>72</sup> | https://github.com/inab/trimal |
| amPEP v1.0 | Lawrence et al.<sup>73</sup> | https://github.com/tlawrence3/amPEPpy |
| ampir 1.1.0 | Fingerhut et al.<sup>74</sup> | https://github.com/Legana/ampir |
| NLPPrecursor | Merwin et al.<sup>36</sup> | https://github.com/magarveylab/NLPPrecursor |
| SignalP 6.0 | Teufel et al.<sup>37</sup> | https://services.healthtech.dtu.dk/services/SignalP-6.0/ |
| peptides.py | N/A | https://peptides.readthedocs.io/en/stable/index.html |
| BAGEL4 | Teufel et al.<sup>37</sup> | https://github.com/annejong/BAGEL4 |
| antiSMASH 6.1.1 | Blin et al.<sup>33</sup> | https://github.com/antismash/antismash |
| MMseqs2 13.45111 | Steinegger et al.<sup>75</sup> | https://github.com/soedinglab/MMseqs2 |
| IQ-TREE version 2.2.0.3 | Minh et al.<sup>76</sup> | http://www.iqtree.org/ |
| GTDB-Tk v2.1.1 | Chaumeil et al.<sup>77</sup> | https://github.com/Ecogenomics/GTDBTk |
| vegan 2.6.8 | Jari Oksanen<sup>78</sup> | https://github.com/vegandevs/vegan |
| iNEXT 3.0.1 | Hsieh et al.<sup>79</sup> | https://github.com/JohnsonHsieh/iNEXT |
| fastp v0.21.1 | Chen et al.<sup>80</sup> | https://github.com/OpenGene/fastp |
| kneaddata | N/A | https://github.com/biobakery/kneaddata |
| PALADIN 1.4.6 | Westbrook et al.<sup>81</sup> | https://github.com/ToniWestbrook/paladin |
| SAMtools 1.16.1 | Li et al.<sup>82</sup> | https://www.htslib.org/ |
| MetaPhlAn version 4.0.6 | Blanco-Míguez et al.<sup>83</sup> | https://github.com/biobakery/MetaPhlAn |
| WGCNA 1.72.1 | Langfelder et al.<sup>84</sup> | https://cran.r-project.org/web/packages/WGCNA/index.html |
| MaAsLin2 1.12.0 | Mallick et al.<sup>56</sup> | https://github.com/biobakery/Maaslin2 |
| ggplot2 v3.4.4 | Villanueva et al.<sup>85</sup> | https://ggplot2.tidyverse.org/ |
| Other |  |  |
| UniRef100 release-2022_05 | The UniProt Consortium<sup>86</sup> | https://www.uniprot.org/help/downloads |
| PFAM version 34.0 | Mistry et al.<sup>87</sup> | http://ftp.ebi.ac.uk/pub/databases/Pfam/releases/Pfam34.0 |
| NCBI version 11.0 | NCBI Resource Coordinators<sup>88</sup> | https://ftp.ncbi.nlm.nih.gov/hmm/11.0/ |
| GTDB repository release 207 | Parks et al.<sup>89</sup> | https://gtdb.ecogenomic.org/downloads |
| GENCODE release 19 (GRCh37.p13) | Frankish et al.<sup>90</sup> | https://www.gencodegenes.org/human/ |

### METHOD DETAILS

#### Development of IIBacFinder

##### Profile hidden Markov models (pHMMs) construction

A total of 286 non-redundant class II bacteriocin sequences (Table S1) were curated from previous studies. To comprehensively identify their homologous sequences, we used jackhammer^69^ to iteratively align them against the UniProt UniRef100 database^86^ (release-2022_05, comprising 323,519,324 sequences) with the parameters “-N 5 -T 20 --domT 20 --incT 20 --incdomT 20”. Putative homologous sequences longer than 150 amino acids were considered false positives and discarded. We manually collated the homologous sequences, removing those with specific functions unrelated to antimicrobial activity or those with unknown functions, resulting in 297,986 remaining sequences. Combined with the 286 known bacteriocin sequences, these class II bacteriocin sequences were deduplicated to avoid significant similarity when constructing pHMMs using CD-HIT with the parameters “-n 2 -p 1 - c 0.8 -d 200 -M 5000 -aL 0.6 -g 1”^68^. The deduplicated sequences were then clustered at a 50% identity threshold using CD-HIT, resulting in 2,884 clusters with low sequence similarity among them. One cluster was removed due to containing a sequence with ambiguous amino acids. We used MAFFT^71^ for multiple sequence alignment within each cluster. Following this, we used trimAl^72^ to automatically remove poorly aligned regions with the model “-automated1”. All aligned sequences were then used to build the pHMM for each cluster using hmmbuild^69^. This process resulted in 2,883 class II bacteriocin-related pHMMs. Additionally, we collected pHMMs from the PFAM^87^ and NCBI databases^88^, obtaining 16 and 14 pHMMs, respectively. To avoid redundancy between the 30 available pHMMs and the 2,883 self-constructed pHMMs, we used hmmscan^69^ to annotate the seed sequences of the 2,883 clusters against the PFAM and NCBI databases. Clusters whose seed sequences were entirely annotated by either PFAM or NCBI domains were discarded, retaining a total of 1,123 clusters. Consequently, we obtained 1,153 precursor-related pHMMs (Table S3).

Subsequently, we curated 29 immunity protein sequences (Table S2) associated with class II bacteriocins from previous reports. To build the pHMMs for these immunity proteins, we first aligned the sequences against the UniRef100 database using Diamond^70^ with the parameter “--id 40 --masking 0”, retaining homologous sequences with identities greater than 40%. These homologous sequences were processed similarly to the precursor sequences, resulting in 29 pHMMs for class II bacteriocin immunity proteins. Additionally, we included eight publicly available pHMMs related to context genes, including protease genes, inducer genes, and accessory genes. Finally, we obtained 37 context gene-related pHMMs (Table S4).

##### Precursor gene identification

IIBacFinder takes a DNA sequence in FASTA format as input, which is subjected to open reading frame (ORF) annotation using Prodigal-short^67^. All ORFs are queried against a library of 1,153 precursor-related pHMMs and 37 context gene-related pHMMs using hmmsearch^69^. Putative precursor genes are detected using both precursor-based and context gene-based rules in parallel. For the precursor-based rule, significant hits identified by the 1,153 precursor-related pHMMs are initially considered putative precursor genes if the hit sequence is less than 150 amino acids in length. The region encompassing ±8 genes around the identified precursor genes is defined as the biosynthetic gene cluster (BGC). For the context gene-based rule, significant hits identified by the 37 context gene-related pHMMs are considered associated with class II bacteriocin biosynthesis. The eight genes upstream and downstream of the identified context genes are included as the putative BGCs. BGC regions lacking a transporter are discarded. Subsequently, the BGC region is examined for the presence of proteins containing peptidase_C39 (PF03412) and LANC_like (PF05147) domains. BGCs containing both domains are discarded as they are likely to be modified lantibiotics. All small ORFs within the BGC, with lengths less than 150 AA, are retrieved and subjected to antimicrobial peptide annotation using both amPEP^73^ and ampir^74^. Small ORFs predicted as antimicrobial peptides (AMPs) with a probability threshold of 0.8 by either tool are considered putative precursor genes. The putative precursor genes detected by either the precursor-based or context gene-based rules are then queried against the Pfam and NCBI databases using hmmscan^69^. Sequences containing domains unrelated to class II bacteriocins are removed (Table S5). The remaining sequences are regarded as putative precursors of class II bacteriocins. Finally, BGC regions detected by both the precursor-based and context gene-based rules are combined.

##### Leader type prediction

The leader type of precursor genes detected by the precursor-based rule is considered identical to that of known homologous bacteriocins (Table S3). For precursor genes detected by the context gene-based rule, we annotate the leader types based on the context genes within the BGC region (Table S4). For example, if the BGC contains an ORF with a peptidase_C39 domain (PF03412), the leader type of the precursor genes is considered to be double-glycine.

##### Leader peptide prediction

To predict the cleavage site of the double-glycine leader, we tuned a deep learning model, NLPPrecursor, specifically designed for the prediction of RiPP precursor cleavage sites^36^. The protein sequences were tokenized using a vocabulary of 24 tokens, including a start and end tag, twenty proteinogenic amino acids, an ambiguous amino acid, and a padding token. Unlike RiPP precursors, which may contain leader and follower sequences at the N-terminal and C-terminal, respectively, double-glycine bacteriocin precursors only have a leader sequence. Therefore, we modified the sequence labels in the model. The allowed transitions of protein sequences within the model were: (start-before), (before-before), (before-propeptide), (propeptide-propeptide), (propeptide-stop), (stop-pad), and (pad-pad). All sequences were unified to a final length of 150 by adding pad tokens at the end.

To train the deep learning model, we created a dataset of double-glycine class II bacteriocins. Since only 142 known class II bacteriocin precursors contain a double-glycine leader, we collected their homologous sequences from the UniProt UniRef100 database to expand the training dataset. DIAMOND^91^ was used to query the 142 sequences against the UniRef100 database with the parameters “--id 50 --query-cover 80 --masking 0”, resulting in 1,224 homologs. To reduce sequence bias in training the prediction model, the 1,336 double-glycine sequences were deduplicated at a sequence identity threshold of 90% using CD-HIT^68^. This deduplication process yielded 439 non-redundant sequences. Subsequently, we used MAFFT^71^ with the parameters “--maxiterate 1000 --genafpair” to conduct multiple sequence alignments and manually check whether the homologous sequences contained a conserved double-glycine cleavage site. Finally, 412 class II bacteriocin precursor sequences with a double-glycine leader were retained for model training with 100 epochs (Figure S1C).

For predicting the Sec-dependent leader, we adopted the well-established tool SignalP 6.0^37^, which predicts the signal peptides of proteins secreted by the secretory (Sec) pathway, using default parameters.

##### Physicochemical property calculation

Following the prediction of the leader peptide, we compute common descriptors for the predicted core peptide using a pure-Python package, peptides.py (https://peptides.readthedocs.io/en/stable/index.html). The physicochemical properties include the aliphatic index, Boman index, isoelectric point, molecular weight, instability index, and net charge at a pH of 7.

##### Comparison with antimicrobial peptides (AMPs)

To gain insight into sequence novelty, predicted core peptides of class II bacteriocins were compared with known AMPs. AMP sequences were retrieved from four publicly available databases in April 2024: APD3^38^, dbAMP^39^, DRAMP^40^, and DBAASP^41^. Sequences less than 10 amino acids or containing ambiguous amino acids were discarded, resulting in 51,692 AMP sequences, of which 21,403 were unique (100% identity). The predicted core peptides were queried against the AMP dataset using DIAMOND ^41^ with the parameters “--ultra-sensitive --max-target-seqs 1 --id 20 --query-cover 20 --masking 0”. The best alignment hit was retained.

##### Prediction confidence assignment

Given that bioinformatic predictions can inevitably yield false positives, we added a confidence label to indicate whether the predicted class II bacteriocins are likely authentic or potentially false (Figure S2C). We considered several factors in determining this confidence level: the presence of a ribosome binding site (RBS), the completeness of the predicted ORF of the precursor gene, the detection rules (i.e., bacteriocin detection based on one rule or both rules), the leader peptide prediction, and the length of the predicted core peptide. The decision tree in Figure S4 illustrates the classification into high-confidence or low-confidence categories. It is important to note that the inferred confidence level does not confirm whether the predicted bacteriocins are authentic or false without experimental validation; it only suggests a likelihood of the prediction’s accuracy.

#### Performance benchmarking on identifying class II bacteriocins

To evaluate the performance of IIBacFinder, BAGEL4^41^, and antiSMASH 6^33^ in detecting class II bacteriocins, we applied these tools to the genomes of known bacteriocin producers. Since most producers’ genomes have not been documented, we first attempted to locate known class II bacteriocins in sequenced bacterial genomes. The ORFs of 257,997 bacterial genomes from the RefSeq database were identified using Prodigal-short^67^ with default parameters. These ORFs were then searched against 286 non-redundant known precursor sequences using MMseqs2 with the parameter “easy-search --max-seqs 1 --min-seq-id 0.95”^75^. A best hit with an identity > 95% and coverage > 95% was considered a known precursor sequence. Consequently, we obtained 137 genomes containing 205 complete and non-redundant precursor sequences of known class II bacteriocins.

We then applied the three prediction tools to these genomes using default parameters to assess their ability to identify class II bacteriocins. Notably, both IIBacFinder and BAGEL4 can pinpoint the precursor sequences, while antiSMASH 6 preferentially detects the biosynthetic region rather than the precursor sequences. Accordingly, if the precursor sequence identified by either IIBacFinder or BAGEL4 was within the BGC region detected by antiSMASH 6, we considered it detected by antiSMASH.

#### Phylogenetic tree construction

The multiple sequence alignment of 286 non-redundant known precursor sequences was achieved by MAFFT v7.508^71^ with the parameter of “--maxiterate 1000 --localpair”, which was further used to infer the phylogenetic tree. IQ-TREE version 2.2.0.3^76^ was adapted for constructing maximum likelihood (ML) phylogenetic trees, with 1,000 ultrafast bootstrap replicates. The best-fit model was chosen as VT+F+I+I+R4 identified by the in-built ModelFinder^92^. The inferred phylogeny was visualized and annotated using iTOL^93^.

#### Analysis of class II bacteriocins in bacterial genomes from multiple datasets

We used IIBacFinder to predict the biosynthetic potential of class II bacteriocins in bacterial genomes from eight datasets, consisting of the RefSeq database (RefSeq, n = 257,997), human microbiome (SGB, n = 126,702), the Unified Human Gastrointestinal Genome (UHGG, n = 285,835), the Human Reference Gut Microbiome (HRGM, n = 29,035), global earth microbiome (GEM, n = 49,466), ocean microbiome (Ocean, n = 21,592), mouse (MGBC, n = 26,640) and ruminant gut microbiome (RGM, n = 10,211). To confirm and unify the taxonomical annotation, GTDB-Tk v2.1.1^77^ was adopted for taxonomical annotation against GTDB-Tk reference data version r207^89^. Genomes not belonging to bacteria were discarded.

#### Rarefaction analysis

To assess the bacteriocin biosynthetic potential of bacterial genomes from the RefSeq database, R package vegan^78^ was used. CD-HIT^68^ was used to group 481,654 high-confidence sequences from RefSeq genomes into 8,610 clusters at a 50% identity threshold The accumulations of bacteriocin clusters detected in Gram-positive and Gram-negative bacteria were computed using *specaccum* function with “random” method.

For the biosynthetic potential in Earth’s microbiome, rarefaction analyses were conducted using the iNEXT R package^79^. We adopted CD-HIT^68^ for deduplicating 10,581 high-confidence sequences from Earth’s microbiome to 4,956 unique sequences at a 100% identity threshold. A unique sequence presence/absence table (sequence versus strain matrix) was constructed for each group considered. The type of input data was set as ‘incidence-raw’ data in the iNEXT main function. Besides, knots of 2,000 and an endpoint of 50,000 were adopted. The number of bootstrap replications is 50 by default.

#### Metagenomics and metatranscriptomics analysis

The raw metagenomic sequencing reads from five cohorts and the raw metatranscriptomic data from three cohorts were acquired from NCBI SRA (Sequence Read Archive) under respective project accession numbers. Only the sequencing data of healthy individuals was retained for analysis. Fastp v0.21.1^80^ with default parameters was used to detect and remove low-quality sequencing reads. The sequencing reads belonging to the human host were detected and discarded by kneaddata (https://github.com/biobakery/kneaddata), through searching against the human reference genome (GRCh38.p13) from GENCODE^90^. The high-quality sequencing reads retained were subjected to downstream analysis.

#### Bacteriocin precursor profiling

Before profiling bacteriocin precursors in metagenomic and metatranscriptomic data, we constructed a reference dataset of class II bacteriocin precursor sequences. IIBacFinder identified 645,318 high-confidence precursor sequences in bacterial genomes from eight databases. To remove duplicates and reduce computational load, we deduplicated these sequences at a threshold of 95% identity and 95% coverage using CD-HIT^68^, resulting in a reference precursor dataset of 32,735 sequences. This reference dataset was then queried by high-quality sequencing reads from metagenomic and metatranscriptomic data using PALADIN software^81^ to profile the abundance of bacteriocin precursors in each sample. We used SAMtools^82^ to calculate their abundance, normalizing the results using transcripts per kilobase million (TPM). To reduce inaccuracies in abundance calculation caused by highly similar sequences in the reference dataset, we grouped the sequences into clusters with a threshold of 80% identity and 80% coverage using CD-HIT^68^, and computed their abundance at the cluster level rather than the individual sequence level.

#### Taxonomic profiling, community diversity, and correlation analysis

Metagenomic taxonomic profiling was conducted using MetaPhlAn version 4.0.6^83^. The relative abundance of species was used to represent the abundance of bacterial species in the community. The alpha diversity (Shannon index) of each sample was calculated with the R package vegan v2.6.4^78^. Given that bioactive bacteriocins could inhibit the growth of other bacteria and thus regulate the overall bacterial community, we computed the Spearman correlation between taxonomic composition (at the species level) or Shannon diversity and precursor clusters using the corAndPvalue function from the R package WGCNA^84^. The p-values were then adjusted using the “BH” method to control the false discovery rate in multiple comparisons^94^. Considering that bioactive bacteriocins would inhibit the growth of specific bacterial species, we retained only the significantly negative associations (*rho* < -0.3 and adjusted *P* < 0.05).

#### Bacteriocin precursor prioritization in metagenomic and metatranscriptomic data

To infer the importance of class II bacteriocins in metagenomic and metatranscriptomic data, we contrived a scheme to assign ranking scores for each profiled precursor clusters, through considering the ecological importance (prevalence and abundance of bacteriocins) and functional importance (association between bacterial species and bacteriocins and between Shannon diversity and bacteriocins). Mathematically, the ecological importance is scored as:

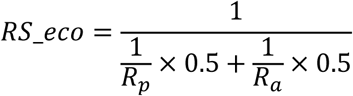

Where RS_eco indicates the ranking score (ecological importance) of a particular bacteriocin within a cohort. Rp and Ra represent the ranking percentiles based on prevalence and mean abundance (largest to smallest), respectively. A bacteriocin with high prevalence and high abundance would give rise to a larger RS_eco.

The functional importance is mathematically scored as:

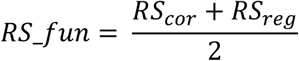

Where RS_fun is the ranking score (functional importance) of a particular bacteriocin within a cohort. RScor indicates the ranking percentiles based on the negative correlation between bacteriocins and bacterial species, while RSreg is the ranking percentiles based on the correlation between bacteriocins and the Shannon diversity index.

The overall ranking score of a bacteriocin in metagenomic and metatranscriptomic data is the average ranking score of ecological and functional importance in all cohorts, mathematically represented as:

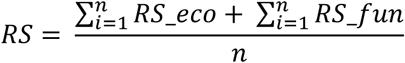

Where n represents the number of metagenomic and metatranscriptomic datasets. A high score of RS implies the great importance of the bacteriocin in the gut microbiome.

After prioritizing the bacteriocins in the human gut microbiome, we selected bacteriocin clusters to validate their antibacterial activities based on the ranking scores (RS) and the following criteria: (1) The hypothetical bacteriocin cluster should include ≥ 5 precursor sequences, as sporadically identified bacteriocins are likely to be false predictions. (2) The cleavage site for double-glycine bacteriocins should be a typical “GG” rather than alternative sites such as “GA” or “GS” to reduce false positives. (3) The length of the deduced core peptide should be less than 50 amino acids due to cost considerations and the feasibility of chemical synthesis. (4) The bacteriocin sequences should not be annotated as peptides with functions other than antimicrobials when queried against the NCBI database. Based on these criteria, we chose to synthesize and test the representative sequences, assigned by CD-HIT^68^, from 40 bacteriocin clusters.

#### Three-dimensional peptide structure prediction

ColabFold was used to predict the three-dimensional structure of peptides, with a parameter of “--amber --templates --num-recycle 3 --use-gpu-relax --model-type auto”^95^. The model with the highest relaxed rank was retained and visualized using PyMol ^96^.

#### Peptide synthesis

The deducted core peptides of predicted bacteriocins were chemically synthesized by solid-phase peptide synthesis by Sangon Biotech (Shanghai, China). Their molecular weights were confirmed by mass spectrometry, and their required purity was greater than 90% determined by high-performance liquid chromatography. The synthesized peptide powder was stored at − 80 °C and dissolved in Dimethyl sulfoxide (DMSO) to 50 mg/mL upon use.

#### Microbial strains and culturing

A total of 33 bacterial strains are included in this study and are detailed in Table S10 regarding the strains and growth conditions. The following media were used: Brain heart infusion medium (BHI); Tryptic Soy Broth with supplement of 0.3% yeast extract (TSB-YE); De Man– Rogosa–Sharpe medium (MRS); Peptone Yeast Glucose medium (PYG); Reinforced Clostridium Medium (RCM); Gifu Anaerobic Broth (GAM); Bifidobacterium Medium (BM); M17 broth. Bacterial strains were grown and streaked on the agar plate, and incubated under respective optimal growth conditions. Individual colonies were then inoculated into 5 mL corresponding culture broth. Once the bacterial culture reached sufficient density, 500 µL culture bacterial suspension was supplemented with 500 µL 50% glycerol to generate liquid glycerol stocks and stored at −80 °C.

#### Antibacterial activity screening

The bacterial inhibition experiment was adopted from a previous study^26^. All bacterial indicators were revived on agar medium under respective growth conditions (Table S10). The individual colonies were picked into broth medium and incubated for one or two days. The bacterial suspension was diluted to a particular concentration [optical density at 600 nm (OD600) = 0.1] and then diluted 100 or 1,000 times for the antibacterial activity screening (Table S10). Three experimental groups were set: (1) blank control group, 100 µL of broth culture; (2) bacterial control group, 50 µL of broth culture and 50 µL of bacterial solution with DMSO (equivalent amount to the solvent of peptides); (3) experimental group, 50 µL of broth culture and 50 µL of bacterial solution with 100 µg/mL peptide. Experiments were conducted on flat bottom 96-well plates. The bacterial growth was measured with OD600 value after statically incubating for 24h or 48h. The inhibition ratio was calculated as:

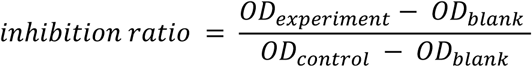

Where ODblank, ODcontrol, and ODexperiment represent the OD600 of the blank control group, bacterial control group, and experimental group, respectively.

All experiments were conducted with three independent replicates. We performed Dunnett’s test for computing the statistical difference between the control and experiment groups. The peptide with a criterion of inhibition ratio ≥ 25% and *P* value < 0.05 was considered effective against the tested indicator strain. We categorized the activity results into four groups: (1) no activity, inhibition ratio < 25% or *P* value ≥ 0.05; (2) low activity, 25% ≤ inhibition ratio < 50% and *P* value < 0.05; (3) medium activity, 50% ≤ inhibition ratio < 75% and *P* value < 0.05; (4) high activity, inhibition ratio ≥ 75% and *P* value < 0.05. Peptides with an inhibition ratio > 90% were further examined the MIC against this indicator strain.

#### Determination of minimum inhibitory concentrations (MICs)

The minimum inhibitory concentrations (MICs) of synthetic peptides were performed by broth microdilution in 96-well flat bottom plates as previously reported^97^. Vancomycin was used as a positive control. Peptides were added to the plate containing 100 μL aliquots of bacterial suspensions and were two-fold serially diluted (ranging from 100 μg/mL to 0.195 μg/mL). After incubating for 24h or 48h, bacterial growth was assessed by determining OD600. The MIC value was defined as the lowest concentration of the peptides where no bacterial or fungal growth was detected. The assays were conducted in triplicate.

#### Human feces collection

Human studies were performed in accordance with all relevant ethical regulations and were approved by the Medical Ethics Committee of ZhuJiang Hospital, Southern Medical University (2022-KY-119). Fecal samples were collected from five healthy Chinese adults aged 20 to 42 who had not taken antibiotics or probiotics within one month before sample collection.

Ex vivo assay was conducted as previously described^17^. Briefly, the collected feces were combined and homogenized in 65 mL of a rich medium mGAM. Following resuspension, the tubes were gently centrifuged at 1,000 rpm for 2 minutes, and the supernatants were then retained for further assay. Four selected bacteriocin peptides were pre-dissolved in DMSO to final concentrations of 0.1 mg/mL, and 10 mg/mL. Subsequently, 10 µL of peptides and 40 µL of fecal supernatant were inoculated into 950 µL of mGAM media, resulting in a final volume of 1 mL. The mixtures were then anaerobically incubated at 37°C for 48 hours. Following the incubation period, the bacterial growth media were centrifuged at 12,000 rpm for 10 minutes. The supernatants were carefully removed, and the resulting pellets were washed twice with 1 mL of PBS before being collected for DNA extraction. The pelleted samples were extracted using the QIAamp® DNA Micro Kit (Qiagen) according to the manufacturer’s instructions. Finally, all DNA samples were prepared for shotgun metagenomics sequencing.

#### Data analysis of human fecal metagenome

All DNA samples were submitted to Novogene for shotgun metagenomics sequencing using a 150bp paired-end protocol as per the manufacturer’s instructions. The raw sequencing data process was described above to obtain the taxonomic profiles. The Vegan package^78^ in R was used to compute alpha diversity and beta diversity at the species level, indicated by Shannon diversity and Bray-Curtis dissimilarity respectively. Differential abundance analysis between groups was conducted using MaAsLin2 (microbiome multivariable associations with linear models) ^56^. Species with a false discovery rate (FDR)-adjusted p-value of less than 0.05 were deemed significantly different between the two groups.

#### Visualization

The intersection of identified bacteriocins from three tools was visualized using the R package UpSetR v1.4.0^98^. The sankey plot showing the biosynthetic potential of bacteriocins in the human gut microbiome was achieved by the package ggsankey v0.0.99999^99^. The heat maps were plotted using the package ComplexHeatmap v2.14.0^100^. The similarity network was visualized using Cytoscape 3.9.0^101^. The organization of gene clusters was visualized by R package gggenes v0.5.0^102^. Without a specific statement, other figures were generated using ggplot2 v3.4.4^85^.

### QUANTIFICATION AND STATISTICAL ANALYSIS

All statistical analyses were finished in R v4.2.3. Dunnett’s test (two sided) was done by the function *DunnettTest* in R package DescTools v0.99.49^103^.

