## Supplementary Figures for "Systematically investigating and identifying bacteriocins in the human gut microbiome"

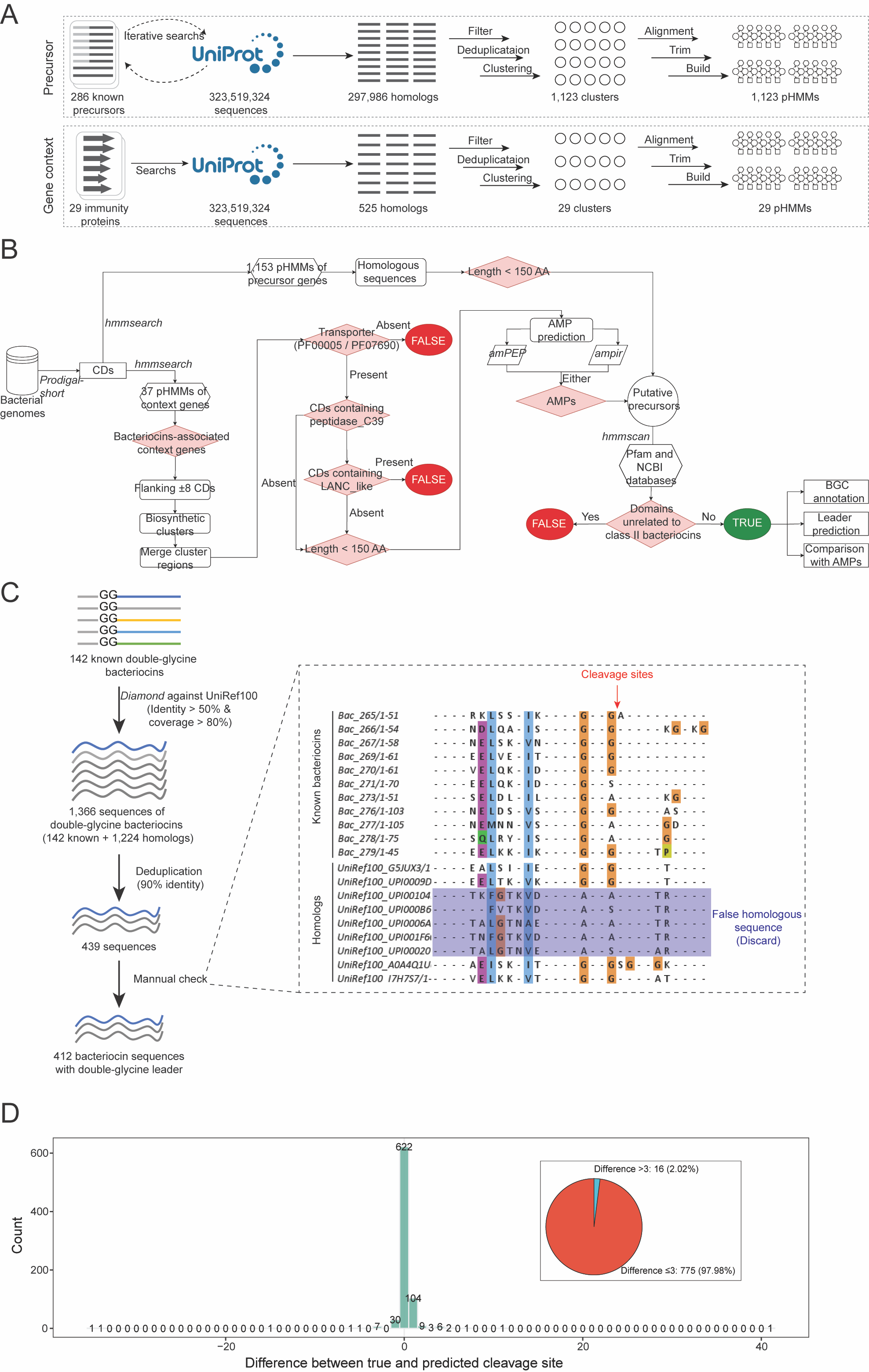

Figure S1. Development of IIBacFinder, related to Figure 1

(A) pHMM construction for precursor and context genes: Initially, 286 non-redundant precursor sequences of known class II bacteriocins are curated from literature. Through iterative searches in the UniRef100 database (323,519,324 sequences), 297,986 homologs are detected and grouped into 1,123 clusters at a 50% identity threshold. Sequences within each cluster are aligned, trimmed, and used to build 1,123 corresponding pHMMs. Similarly, for context genes, 29 distinct genes responsible for conferring self-immunity to class II bacteriocins are curated and used to build 29 pHMMs.

(B) Schematic flow of bacteriocin mining in IIBacFinder.

(C) Curation of class II bacteriocins with double-glycine leaders: Initially, 142 known class II bacteriocin precursors with double-glycine leaders are aligned against the UniProt UniRef100 database to identify homologous sequences, resulting in 1,336 double-glycine sequences. To reduce sequence bias in the prediction model, these sequences are deduplicated at a 90% identity threshold, leaving 439 non-redundant sequences. After multiple sequence alignment, 28 sequences deemed potential false predictions are discarded. Ultimately, 412 class II bacteriocin precursor sequences with double-glycine leaders are retained for model training.

(D) Bar plot showing the distribution of differences between true and predicted cleavage sites, computed from 10-fold cross-validation. The pie chart depicts the percentage of differences less than 3 amino acids.

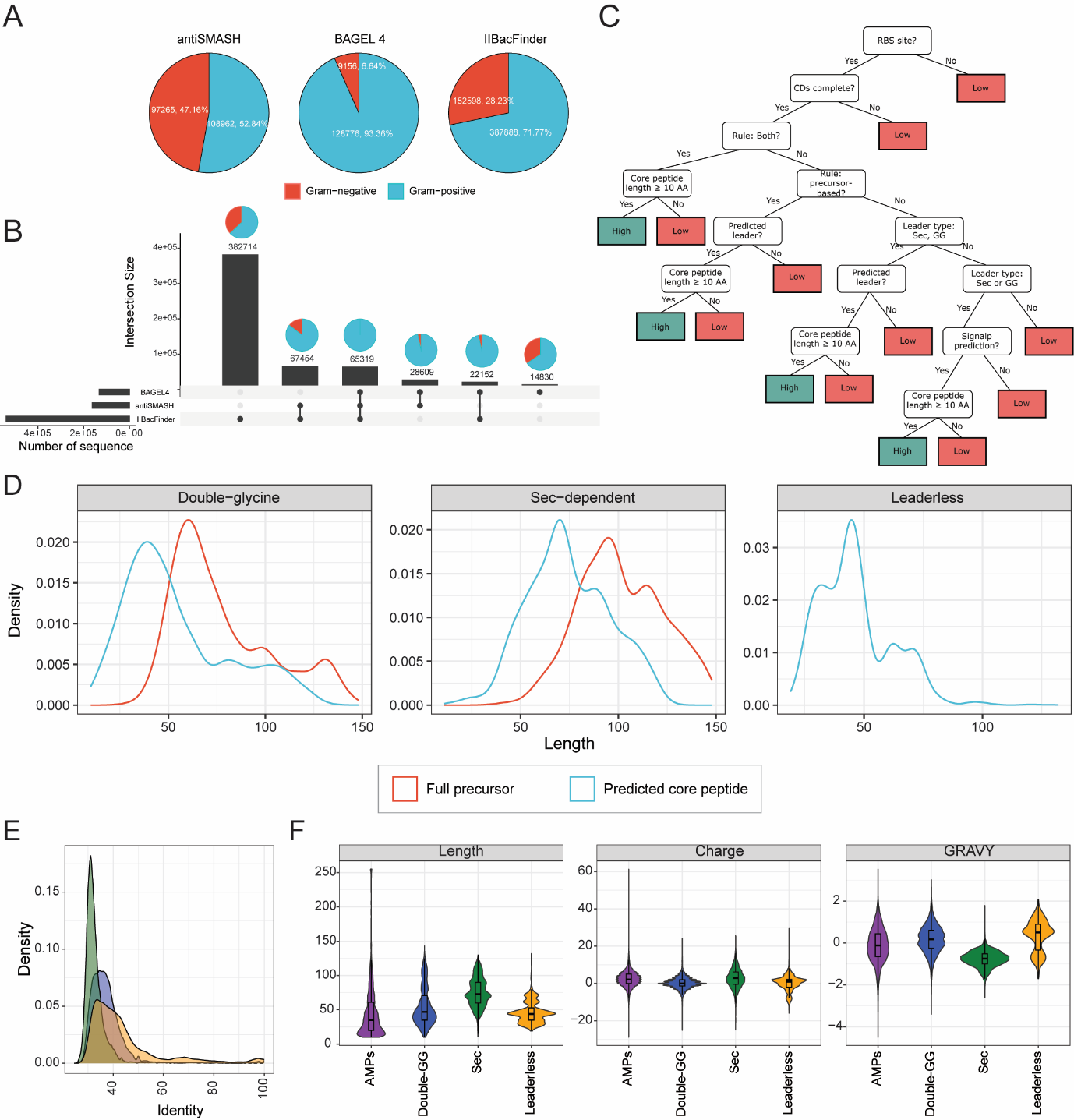

Figure S2. Identification of Class II bacteriocins from RefSeq bacterial genomes using three prediction tools, related to Figure 2

(A) Pie charts showing the proportion of identified bacteriocin sequences from Gram-positive and Gram-negative bacteria, using three prediction tools (i.e., antiSMASH 6, BAGEL 4, and IIBacFinder).

(B) The bar plot on the left refers to the number of bacteriocin sequences identified by three tools. The bar plot on the top depicts the number of bacteriocin sequences of each intersection. Connecting lines are drawn if an intersection is present in more than one tool. The pie charts show the proportion of bacteriocin sequences from Gram-negative and Gram-positive bacteria in each intersection.

(C) Decision tree for classifying the prediction confidence. The predicted bacteriocin sequence is first examined for the presence of ribosome binding site (RBS) and CDs completeness inferred by the ORF calling tool (i.e., Prodigal-short). For further classification, three criteria are included, including the detection rule (precursor-based rule or gene-context-based rule), leader prediction, and length of the deduced core peptide. Additionally, to ensure accuracy, a criterion is included that eliminates any putative core peptide less than 10 amino acids in length, as no Class II bacteriocins have been found to be less than 10 amino acids in length. Through manual scrutiny, it has been determined that sequences with a putative core peptide length less than 10 amino acids are more likely to be false predictions.

(D) Density plots illustrating the distribution of the length of full precursor sequences and deduced core peptides.

(E) Density plot showing the distribution of identity between AMPs and three types of bacteriocins (double-glycine, sec-dependent, and leaderless).

(F) Violin plots depicting the distribution of length of AMPs and core peptides of bacteriocins (left panel), charge at pH = 7.0 (middle panel), and GRAVY (right panel). GRAVY index scores represent the average hydrophobicity and hydrophilicity. The figure legends are shared between figures (E) and (F).

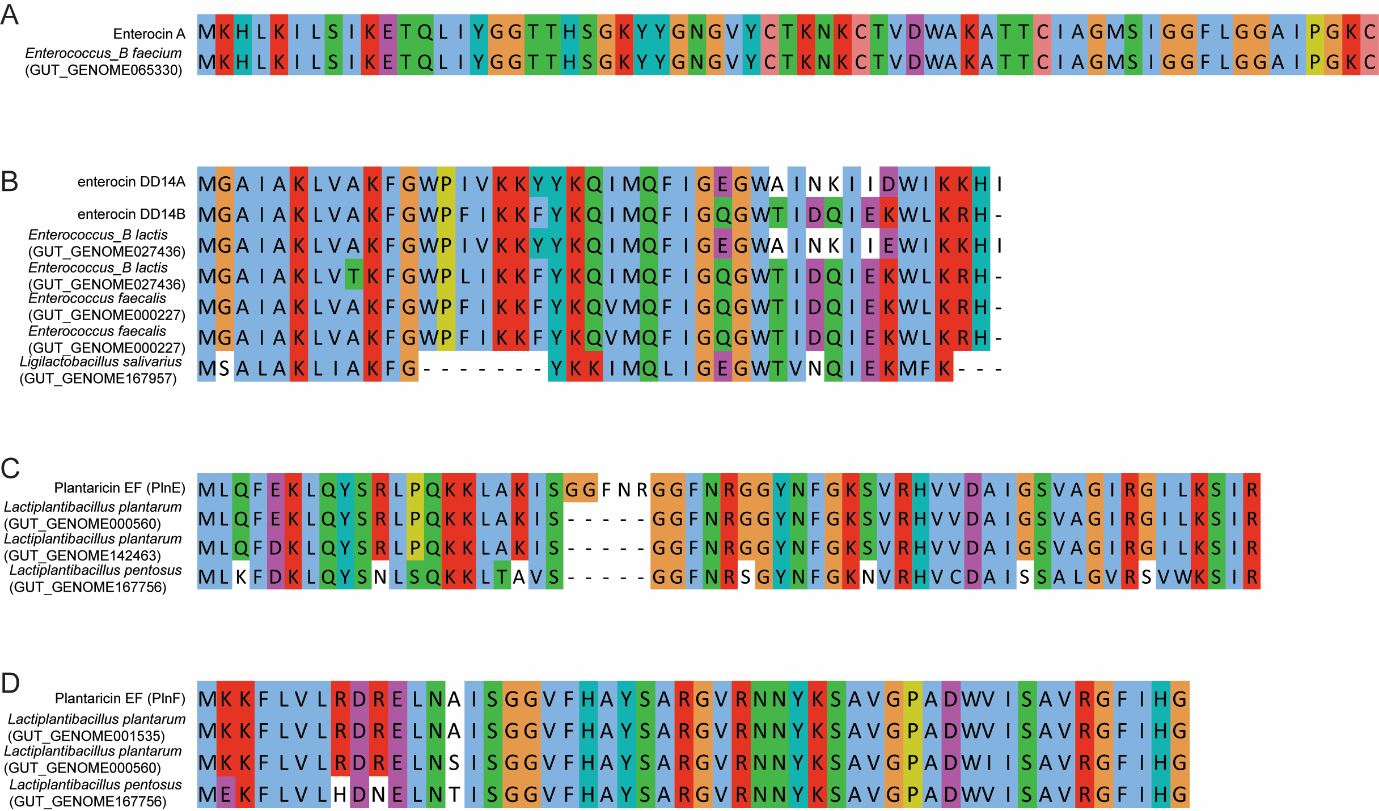

Figure S3. Human gut microbiota-derived class II bacteriocins homologous to functionally known bacteriocins, related to Figure 4

(A) Homologous sequences of Enterocin A.

(B) Homologous sequences of enterocin DD14.

(C) Homologous sequences of Plantaricin EF PlanE.

(D) Homologous sequences of Plantaricin EF PlanF. Genomes harbouring corresponding bacteriocin genes are indicated in the brackets.

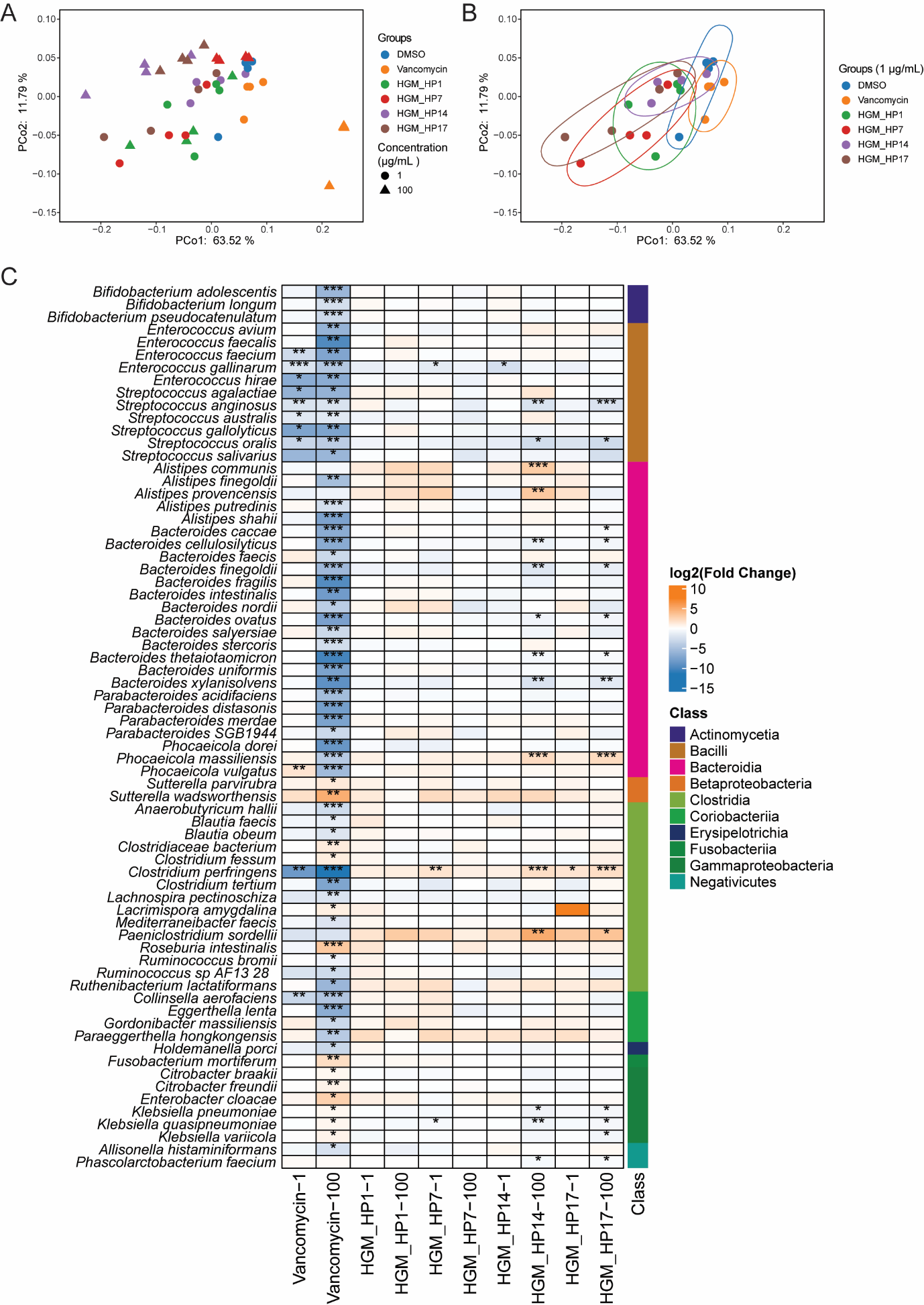

Figure S4. Changes in microbial communities with treatment of antibiotic and bacteriocins, related to Figure 6

(A) Principal Coordinate Analysis (PCoA) of the microbial community of all groups, based on Bray-Curtis dissimilarity at the species level.

(B) Principal Coordinate Analysis (PCoA) of the microbial community of groups with the treatment of 1 µg/mL of vancomycin or bacteriocins, based on Bray-Curtis dissimilarity at the species level.

(C) Differentially abundant species. The heatmap shows the significantly differentially abundant species between treatment groups vs. the DMSO group. Each treatment group is symbolized by the bacteriocin and the concentration tested (e.g., HGM_HP1-1 represents 1 µg/mL of HGM_HP1, whereas HGM_HP1-100 represents 100 µg/mL of HGM_HP1). Statistical significance is denoted as follows: **P* < 0.05; ***P* < 0.01; ****P* < 0.001; ns, not significant.
